# The influence of parental and genotype effects on early survival and development in Atlantic salmon

**DOI:** 10.64898/2026.08.14.744584

**Authors:** Katja S. Maamela, Jenni M. Prokkola, Corinne Suvanto, Xin-Di Huang, Craig R. Primmer, Kenyon B. Mobley

**Author notes:** Corresponding authors: Katja S. Maamela,; Kenyon B. Mobley. These authors share senior authorship (Craig R. Primmer, Kenyon B. Mobley).

## Abstract

Parental qualities can influence the development and fitness of their offspring via genetic and non-genetic effects. Although these effects are often linked to parental phenotypes, the effect of parental genetic variation linked with relevant phenotypes is less well understood. We performed full factorial crosses based on parental genotypes for an age-at-maturity-related gene, *vgll3,* to investigate how the parental genotypes influence Atlantic salmon (*Salmo salar*) offspring survival, growth, and development in their early life. Beyond the connection with age at maturity, the additional association between *vgll3* and body condition in Atlantic salmon offers a potential pathway by which the maternal *vgll3* genotype could influence offspring early life fitness. Combined with measurements of maternal phenotype and egg characteristics, the crossing design therefore allowed us to disentangle the maternal and paternal genetic and non-genetic contributions to variation in offspring survival and phenotypic traits. The phenotypic traits measured were hatching length and yolk sac area, growth, and yolk sac consumption and conversion efficiency. Parental *vgll3* genotype did not influence the majority of our measured egg traits or alevin traits except for a genetic effect of paternal *vgll3* genotype on offspring survival, whereby the paternal late maturation allele was associated with higher survival. Maternal effects were strongest for survival and for traits associated with hatching and weaker for alevin growth and yolk sac usage. Paternal effects on the measured alevin traits were negligible. The results from our study demonstrate that both maternal and paternal effects have the potential to influence offspring early life fitness traits.

## Introduction

Parental effects play an important role in offspring survival and development in a wide range of species (Uller, 2012). Parental effects can be defined as the causal influence of parental phenotype or genotype on offspring phenotype that is beyond the effect of inherited genes (Crean & Bonduriansky, 2014; Mousseau & Fox, 1998; Wolf & Wade, 2009). For example, parental effects can manifest as maternal provisioning of nutrients and hormones (Bernardo, 1996b; Räsänen et al., 2005; Sharda et al., 2021; Uller, 2012), environmental influence on offspring traits due to parental choice of nesting site (Lloyd & Martin, 2004) or immune factor transfer to offspring (McNamara et al., 2014; Saino et al., 2002). Despite our knowledge of how such parental effects potentially influence offspring fitness traits (Burton et al., 2020; Giesing et al., 2011; Mitchell et al., 2013; Warner & Lovern, 2014), disentangling the contributions of parental genetic and non-genetic effects would help clarify their relative contributions to offspring fitness. Utilizing prior knowledge of the genetic basis of reproduction-linked life-history trait variation (Ayllon et al., 2015; Barson et al., 2015; Küpper et al., 2016; Lamichhaney et al., 2016) allows the study of genetic and non-genetic parental effects on offspring fitness traits.

Of parental effects, maternal effects have received more attention compared to paternal effects especially in species with little to no parental care. This difference in knowledge is likely due to the large contribution of the female via maternal provisioning of eggs while the male’s role may be limited to sperm and seminal fluid transfer (Bernardo, 1996a; Crean & Bonduriansky, 2014; Mousseau & Fox, 1998). Though paternal effects have traditionally received less attention, they are now being recognized as an important contributor to offspring phenotypic variation (Crean & Bonduriansky, 2014; J. P. Evans et al., 2019; Rutkowska et al., 2020). For example, in a fish species that has paternal care, antimicrobial compounds may be provided to offspring to help stave off infections (Giacomello et al., 2006). Taken together, both maternal and paternal effects may provide benefits to offspring fitness and contribute significantly to phenotypic variation (Crean & Bonduriansky, 2014).

Atlantic salmon (*Salmo salar*) offers a great opportunity to study parental effects as their eggs are large with a long developmental time that can be manipulated with water temperature (Gunnes, 1979) and the eggs and hatching larvae can be grown in controlled common-garden settings to disentangle environmental and genetic effects on offspring phenotypic traits. Additionally, Atlantic salmon do not provide extensive parental care to their offspring, which simplifies the partitioning of parental effects on offspring fitness (Neff & Pitcher, 2005). During spawning, females choose spawning sites and excavate nests (called redds) in the river substrate. Spawning may take place over several days with one or more males after which the redd is covered with gravel. Both females and males leave the redd after spawning and no parental care is given beyond the female choice of the redd location and construction (Fleming & Einum, 2011).

Maternal effects in the form of egg provisioning is an important means by which a mother can influence the fitness of their offspring (Berg et al., 2001; Bernardo, 1996b; Brooks et al., 1997; Heinimaa & Heinimaa, 2004; Mousseau & Fox, 1998). In Atlantic salmon, ovarian maturation and provisioning of the eggs with essential components (Brooks et al., 1997) takes place during a period prior to spawning when the females do not feed and consequently, the females use their somatic reserves for egg provisioning (Kadri et al., 1996). This use of acquired somatic reserves highlights the importance of maternal resource acquisition and allocation on egg composition as higher maternal energy reserves may be associated with higher egg energy provisioning (Ouellet, 2001). Due to lack of extensive parental care in Atlantic salmon, maternal provisioning via egg yolk and its energy content becomes a crucial resource for offspring survival and development (Berg et al., 2001). The yolk is an essential source of energy for the embryo and hatched larvae (Kamler, 2008). The alevins (i.e., the yolk-sac-feeding life stage after hatching) develop with yolk lipids and proteins serving as the main sources of energy prior to independent feeding (Berg et al., 2001; Brooks et al., 1997) underscoring the importance of maternal provisioning of her offspring. It is unclear to what extent paternal effects exist in Atlantic salmon as the male’s contributions to reproduction are sperm and seminal fluids.

In Atlantic salmon, age at maturity (i.e., the age at which an individual returns to freshwater to spawn for the first time) is mediated by environmental and genetic factors (Mobley et al., 2021). Atlantic salmon can spend anywhere from one to multiple winters at sea prior to maturation (Fleming, 1996; Mobley et al., 2021). A large-effect locus, *vgll3,* explains ∼39% of the variation seen in age at maturity in wild Atlantic salmon, where the E allele is associated with a higher probability to mature at an earlier age and the L allele at a later age, respectively (Barson et al., 2015). This association between *vgll3* and age at maturity has thereafter been demonstrated in common-garden studies using both male (Åsheim et al., 2023; Ayllon et al., 2019; Debes et al., 2021) and female (Maamela et al., 2025) salmon. However, this effect of *vgll3* on age at maturity in Atlantic salmon is not directly related to growth (Debes et al., 2021).

The *vgll3* locus is conserved in vertebrates and has been linked to several maturation-related processes such as pubertal timing in humans (Cousminer et al., 2013) and adipogenesis in mice (Halperin et al., 2013), and sex-biased diseases in humans (Liang et al., 2017; Plazyo et al., 2025). In Atlantic salmon, previous studies have found an association between *vgll3* and body condition (Debes et al., 2021; House et al., 2023), further linking *vgll3* with the energetics of maturation. Body condition is a proxy for available somatic energy reserves (e.g., lipids) that can be allocated to reproduction (Shearer & Swanson, 2000) and higher body condition is associated with higher probability of maturation across taxa (Bernardo, 1993). For female salmon, age at maturity is also associated with offspring fitness via the correlation between female body size and egg size (Fleming, 1996). Larger body size (and older age) at reproduction is positively correlated with not only fecundity, but also with egg size and energetic resource allocation of the eggs (Heinimaa & Heinimaa, 2004). Thus, maternal *vgll3* genotypes could influence egg provisioning via effects on body condition and size at maturity providing a potential pathway by which *vgll3* could influence offspring fitness.

The role and phenotypic effects of *vgll3* during early life-history (e.g., post-hatch growth and development) is not well established. However, previous studies have shown that five-month-old Atlantic salmon juveniles with differing *vgll3* genotypes vary in levels of aggressive and exploratory behaviour (Bangura et al., 2022, 2024), and at the molecular level, *vgll3* is expressed in embryo and alevin (i.e., hatched larva with egg sac still attached) stages (Kurko et al., 2020). Further, juveniles with different *vgll3* genotypes show varying gene expression patterns related to lipid storage regardless of maturation status (Ahi et al., 2025). Additionally, the brain-pituitary-gonad axis genes are differentially expressed between *vgll3* genotypes in multiple tissues that may affect age at maturity and reproduction (Ahi et al., 2022). These gene expression differences may lead to offspring fitness effects if the genetic differences are associated with parental gamete quality. Results from a recent study hint at a link between *vgll3* and offspring survival where parental late maturation *vgll3* genotypes were associated with higher offspring early-life survival in wild Atlantic salmon compared to offspring of the early maturation *vgll3* genotype parents (Aykanat et al., 2024). How offspring growth and development may be influenced by parental *vgll3* genotypes in combination with different maternal phenotypes and egg traits has yet to be explored.

In this study, we investigated how parental effects and parental genotypes of the maturation-related gene *vgll3* influence offspring fitness traits in Atlantic salmon in their early life. Specifically, our first goal was to investigate whether female phenotype and *vgll3* genotype influenced egg traits such as egg size and egg lipid and protein contents. We expect that larger females will invest more into eggs (Heinimaa & Heinimaa, 2004) that confer higher fitness benefits to developing offspring. Our second goal was to investigate how parental effects and parental *vgll3* genotypes affect survival up to the first-feeding stage. We predict that survival will be dependent on parental effects more than parental *vgll3* genotype due to the strong potential for egg provisioning and gamete quality to influence offspring survival (Einum & Fleming, 1999). Our third goal was to investigate how egg traits, parental *vgll3* genotypes, and overall parental effects influence alevin body and yolk sac sizes at hatching as well as growth and yolk sac usage. We predict that alevin hatching phenotype, growth, and yolk sac usage will be more influenced by maternal effects than paternal effects or parental *vgll3* genotypes due to the observed influence of maternal effects on offspring early-life in salmonids (Burton et al., 2013; Falica et al., 2017; Houde et al., 2015; Leblanc et al., 2016). Finally, we wanted to compare the overall maternal and paternal effects on offspring early-life traits. Similar to our third goal, we expect maternal effects to be larger than paternal effects due to the high maternal investment into eggs. To address these goals, we raised families of Atlantic salmon crossed based on parental *vgll3* genotypes in a full-factorial crossing design. In this manner, we could partition the individual effects of maternal and paternal *vgll3* genotype as well as estimate the non-genetic contribution on survival, egg traits, and offspring developmental traits.

## Methods

### Study animals

The parental individuals used in this study were from first-generation hatchery broodstock originating from the river Iijoki (65.32°N, 25.43°E) in Finland, reared at Natural Resources Institute Finland (Luke) Taivalkoski hatchery. The hatchery strain was first founded in the 1960s after hydropower dams blocked migration to spawning grounds and new cohorts have been created with returning spawners every few years since then. The broodstock used in this study was created in 2017 when 18 female and 18 male salmon caught on their return migration to the river Iijoki were crossed by dividing each female’s egg clutch into five parts and fertilizing each part with milt from five different males. The broodstock was then used for the first time for supplementary stocking in autumn 2022. Individual-level information regarding maturation and previous spawning events are not available for this broodstock. However, based on the broodstock age, it can be estimated that parental individuals used for this study were spawning either for the first or second time in autumn 2023 when gametes were collected for this study.

### Pedigree reconstruction

In 2022, the broodstock was PIT tagged and finclipped for genetic analyses. The genotypes of the individuals were analysed using a 177 single nucleotide polymorphism (SNP) panel (Aykanat et al., 2016) sequenced on an Illumina platform (Next-Seq). Relatedness among the individuals was analysed using a likelihood ratio method and these relatedness results were then used to construct a pedigree via the R package *SEQUOIA* v2.11.2 (Huisman, 2017). This pedigree was used to select unrelated parental fish for the experimental crosses.

In October 2023, eggs and milt were collected from selected parental fish representing all *vgll3* genotypes at the Luke Taivalkoski hatchery (65.60°N, 28.08°E) to enable crosses between unrelated individuals with a variety of *vgll3* genotype combinations. We sampled eggs of 39 mature females (13 EE, 11 EL, and 14 LL) and milt of 39 mature males (11 EE, 14 EL, and 14 LL males). At stripping, fork length (mm) and wet mass (kg) (both pre- and post-stripping the eggs) of the females were recorded. Fork length is used as the measure of maternal body length throughout this study. Post-stripping wet mass and fork length were used to calculate body condition for the females as Fulton’s condition factor with the formula K = 100 x (mass (g)/fork length^3^ (cm)) (Heincke, 1908).

### Maternal investment traits

All egg traits were measured from unfertilized eggs and represent within-female averages. Egg size was measured as both the average egg wet weight and dry weight. A day after stripping, egg wet weight was measured by weighing three replicates of 10 unfertilized eggs (in g) each and calculating the single egg weight as mean of these three replicates. Egg dry weight was measured by drying a sample of 10 eggs (stored in -80°C freezer until drying) for 48 hours at 60°C and calculating the mean individual egg dry weight from this sample.

Egg total lipid content was quantified using the lipid extraction method based on Folch et al. (1957). Briefly, a pre-weighed sample of 10 eggs was split into two replicates of 5 eggs each. The eggs were first homogenized in 900 μL of milliQ water using an Omni Bead Ruptor Elite (Omni International) and the homogenate was transferred to a pre-weighed 15 mL glass vial. Lipids were extracted into chloroform by adding 9mL of chloroform:methanol solution (2:1 vol/vol) into the homogenate. After extraction, the solvent was evaporated under a constant nitrogen stream and the remaining lipid residue was weighed (to 0.0001 g). The mean single egg lipid content (in mg) was calculated as the average weight of lipids from the two 5-egg replicates.

Egg protein content was quantified using the Bradford method (Bradford, 1976) following the microplate protocol of the Pierce™ Coomassie (Bradford) Protein Assay Kit (ThermoScientific, Rockford, IL, USA). A pre-weighed sample of 10 frozen eggs was dissolved in 4 mL of NaOH (0.1 M) in a 60°C water bath. Due to the high protein content of the eggs, the protein subsamples were diluted (1:200) with NaOH (0.1 M) before the analysis. The protein concentration of the sample was measured using a spectrophotometer (EnSpire Multimode Plate Reader, PerkinElmer) at 595 nm. The average protein concentration of an individual egg (mg * mL^-1^) was calculated as the average of two samples. Standardization of protein concentration was accomplished using a linear curve of a serial dilution of bovine serum albumin.

### Crossing design

Eggs and milt were transported on ice overnight to the Viikki campus of the University of Helsinki where fertilizations were conducted the day after collection. To study the potential influence of maternal and paternal *vgll3* genotype on alevin early-life survival and growth, unrelated parental individuals were crossed in eleven complete 3 x 3 factorials resulting in 99 families in total (Figure 1). In total, the crosses included 33 females (11 females of each *vgll3* genotype) and 31 males (11 EL and LL males, 9 EE males) of the sampled adult salmon. Three males, all EE, were excluded from the crosses due to unviable sperm (i.e., no movement when activated with water). Therefore, two of the EE males were used in two separate factorials. Approximately 200 eggs were used in each cross.

**Figure 1.**
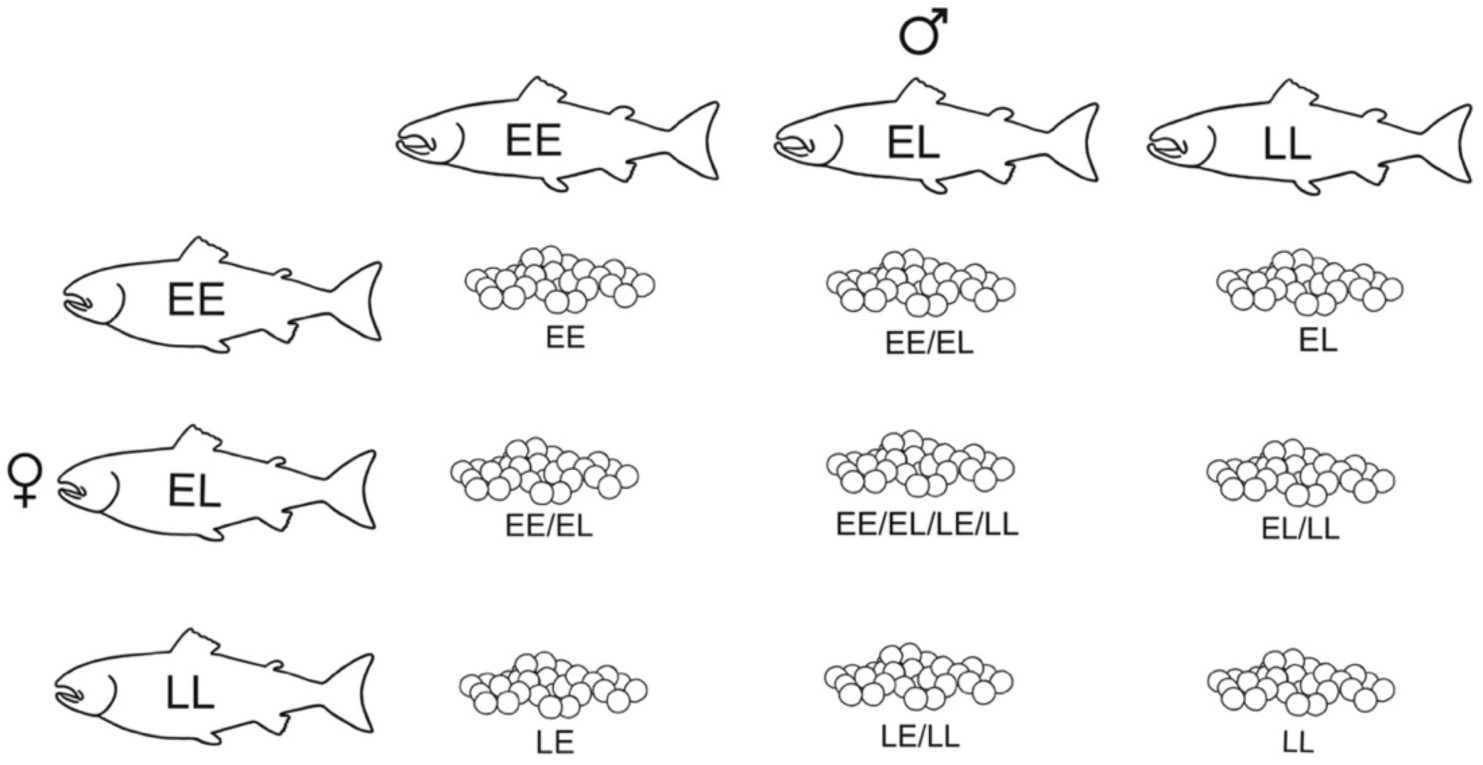
Experimental crossing design. Eggs and milt were crossed in eleven 3 x 3 factorials based on parental *vgll3* genotype. In total, the crosses included 33 females and 31 males. Genotypes under egg images indicate the expected genotypes observed in offspring of a particular cross and the allele(s) inherited from the mother is listed first.

### Family-level egg rearing and survival

Eggs were incubated at 7.5°C in two vertical recirculating incubators (MariSource Inc., WA, USA) housed in a dark experimental room with each family split into roughly two equal sized replicates and assigned to trays in the incubators randomly using a random number generator. Unfertilized eggs (i.e., eggs that were completely white) were removed from the incubators 24 hours after fertilizations. These removed eggs were not counted. Key developmental timepoints for the embryos and alevins were estimated at this temperature using tau (t_s_) units (Gorodilov, 1996). The calculated t_s_ units indicate the time that an embryo takes to form a somite pair at a specific temperature and can be used to estimate the expected date of crucial developmental timepoints (e.g., eyed-stage, hatching, and first feeding) given the water temperature (Gorodilov, 1996). Water temperature was monitored on each incubator shelf using temperature loggers (HOBO TidbiT MX2203, Onset Computer Corporation, MA, USA). Water temperature did not differ between the two incubators (Figure S1) and the average water temperature from fertilization until first-feeding stage was 7.32 ± 0.003 °C. On 14 January 2024 (93 dpf), there was a water temperature control system failure in the incubators, which caused a drop in water temperature down to approximately 4.5°C for one day. No obvious effect on alevin survival due to the temperature drop was observed.

Dead eggs were removed from the incubators after egg shocking at the eyed stage and counted. After hatching had commenced and lasting until first feeding (109 days post fertilization; dpf), the incubators were checked several times for dead alevins, which were then removed and counted. Checking the incubators for dead alevins was infrequent to avoid unnecessary stress for the alevins. At first feeding when the experiment ended, the number of dead and alive alevins was recorded for each family (family replicates pooled). Family-level survival was then calculated as the number of alive alevins surviving to first feeding out of the total number of alevins and eggs (dead and alive) per family. Alevins which were raised in the tissue culture plates (see below) were removed from the total alevin count of their respective families prior to calculating family-level alevin survival.

### Individual-level alevin rearing

At the eyed egg stage (41 dpf, approximately 170 t_s_), individual eyed eggs from a sub-sample of families were randomly selected and placed in wells filled with 10 mL of sterile oxygenated water on 6-well microtiter plates (randomized) to allow for individual-level monitoring of growth. In total, eyed eggs were sampled from 72 families from eight complete 3 x 3 factorial crosses. We excluded three 3 x 3 matrices due to low survival in families of specific females (n = 2) or due to one female being related to another female used in the crosses (n = 1) based on pedigree information. The number of eggs sampled from each family depended on the cross type of the specific family as follows: 12 eggs were sampled from families in crosses where both parents were *vgll3* homozygotes or *vgll3* homozygote-heterozygote crosses, and 24 eggs were sampled from crosses where both parents were heterozygotes. In total, this resulted in 960 eggs being randomly allocated onto 160 6-well microtiter plates with lids (total well volume 16.8 mL, Corning Costar). Water level in the wells was monitored throughout the experiment and new sterile oxygenated water was added when evaporation occurred. The well plates were distributed onto 10 shelves in two climate chambers (Climacell, MMM group, München, Germany) with temperature set to 7.5°C in the dark.

### Alevin traits

Hatching started at 55 dpf (∼258 t_s_) with peak hatching occurring at 64 – 66 dpf (∼300 – 309 t_s_). Alevins were checked daily starting from when the first alevin hatched and each day, plates with newly hatched alevins were photographed to obtain body length and yolk sac size measurements at hatching. After all alevins completed hatching (68 dpf), the alevins were photographed weekly for measurements of body length and yolk sac area. To collect these phenotypic measurements, the well plate was placed on top of a continuous light source and a photograph was taken using a digital camera (Nikon D300s fitted with an AF-S Micro Nikkor 105mm f/2.8G IF-ED macro lens, aperture f/11, shutter speed 1/80s) mounted on a tripod directly above the well plate. The position of the tripod and camera remained the same through the experiment and the plates were placed on roughly the same location on the light table during photographing. A ruler was included in the photos for scale. Before photographing, the wells were filled to the top with water (sterile and oxygenated) in order to standardize the amount of magnification/distortion of alevins caused by water in the wells. Alevins were not anaesthetized prior to photographing. Therefore, to minimize the distortion caused by water at the edges of the wells (see e.g., Figure 2 well 1), a photograph was taken after the alevins had stopped moving and alevins close to the edge on the well were gently repositioned towards the well centre using the tip of a plastic Pasteur pipette.

**Figure 2.**
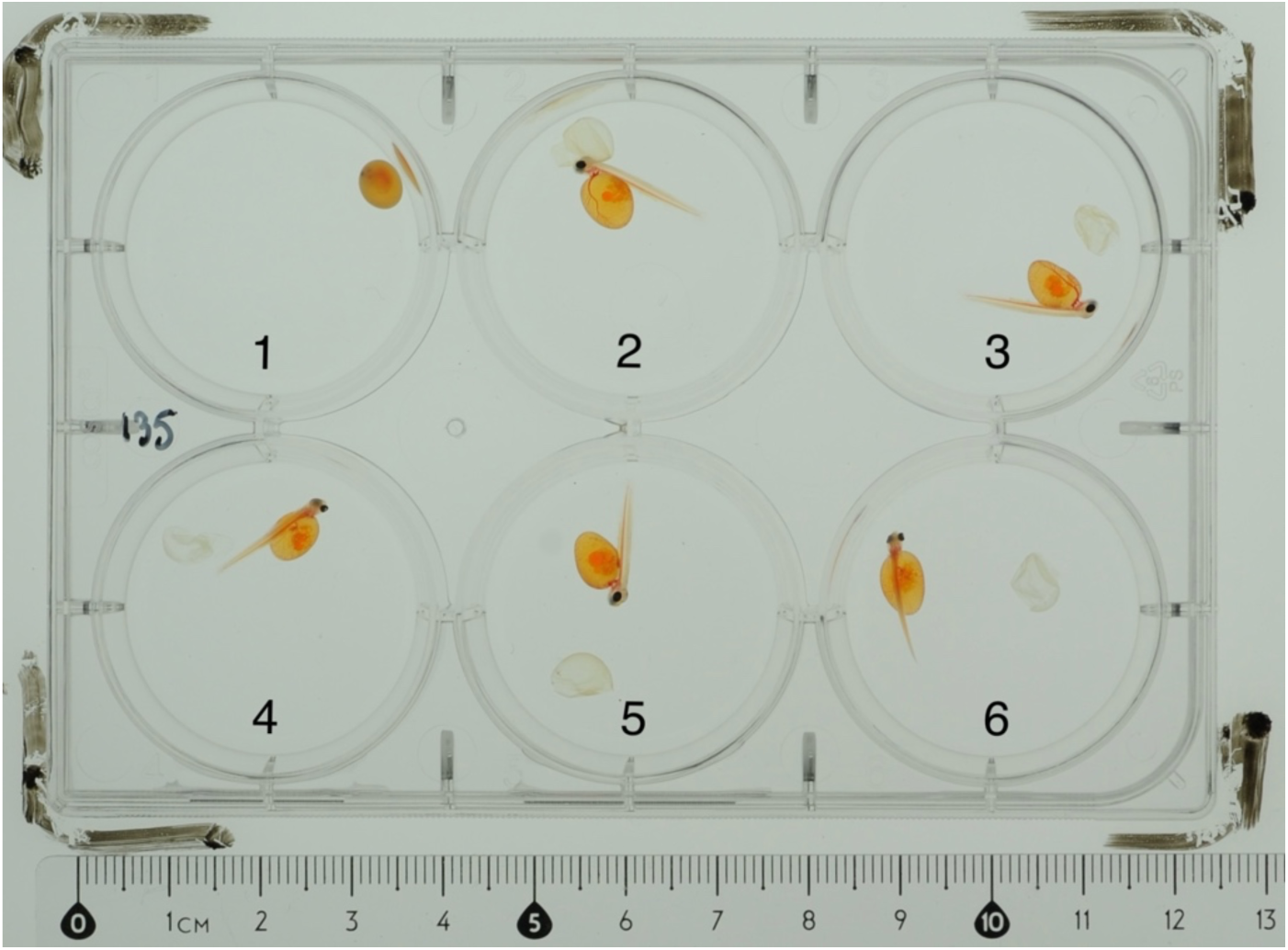
A photograph of hatched alevins on a well plate. Alevins were categorized based on their body position into one of four categories outlined in Table 1. Alevins in wells 2, 3, and 5 belong in category 1, alevin in well 4 in category 2, and alevin in well 6 in category 3. Individual in well 1 had not hatched.

**Table 1.** Description of the photograph quality criteria used to assign photographs into categories which were used to exclude less accurate measurements.

| Category | Criteria |
| --- | --- |
| 1 | Lateral side of the alevin fully visible. Lateral view of the yolk sac fully visible. Both alevin body length and yolk sac area measurements available. |
| 2 | Alevin laying partially on top of its yolk sac and lateral side partially visible. Lateral view of yolk sac partially visible. No yolk sac area measurement available. |
| 3 | Dorsal view of the alevin visible. Alevin laying on top of its yolk sac. No yolk sac area measurement available. |
| 4 | Alevin moving in the photograph. No alevin body length or yolk sac area measurement available. |

The quality of phenotypic measurements was assessed using four different categories based on alevin body position in the photograph (Table 1). Only measurements from photographs belonging to category 1 were included in the final analysis. Alevin body length (mm) and yolk sac area (mm^2^) were measured from the photos using a macro in ImageJ version 1.53s (Rueden et al., 2017). Alevin body length was measured as the length of a spline from the tip of the snout along the visible caudal artery until the base of the caudal fin. As the caudal fin is not clearly visible on all the photographs, the fin was not included in the length measurements. Yolk sac area was measured as the area of a polygon traced along the edge of the yolk sac.

We measured alevin developmental traits (i.e., growth and yolk sac usage) between two timepoints; at hatching and at 81 dpf. The second timepoint at 81 dpf was chosen as it was the latest development point prior to signs of water quality issues (i.e., elevated ammonium and nitrite concentrations) becoming evident that may have influenced growth and development. In all calculations, time is measured as dpf.

Alevin growth rate (G) was calculated using the formula:

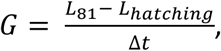

where *L* is alevin body length (mm) and Δ*t* is the number of days between hatching and phenotype measurements at 81 dpf.

In addition to growth, we calculated rate of yolk sac consumption from hatching until 81 dpf as a proxy for energy resource use. Yolk sac consumption rate (YCR) was calculated as:

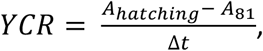

where *A* is yolk sac area (mm^2^) and Δ*t* is number of days between hatching and phenotype measurements at dpf 81.

To investigate how the alevins are converting their yolk sac into growth, we calculated yolk sac conversion efficiency (YCE). YCE was calculated using the formula:

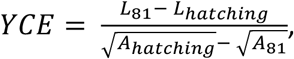

where *L* is alevin body length (mm) and A is yolk sac area (mm^2^). Thus, YCE is a ratio between the change in body length and change in yolk sac area between hatching and 81 dpf.

### Statistical methods

#### Samples used in the models

Of the 39 sampled females, 38 were used in the fecundity and egg trait models after removing one female with missing phenotypic data. Of the 960 eggs that were sampled onto the well plates, 473 alevins were included in the analysis of body length and yolk sac size at hatching after removing individuals that did not hatch, were deformed, had maternal phenotype or egg trait data missing, or did not have an accurate measurement of alevin body length or yolk sac area (measurements classified in photo quality category 2, 3 or 4). The models for growth, yolk sac consumption rate, and yolk sac conversion efficiency included 285 alevins after only including alevins that had accurate phenotypic measurements for both timepoints and removing alevins that died between hatching and the second phenotypic measurements at 81 dpf. For the summary statistics of maternal and alevin phenotypic traits, we report the means and one standard error of the mean (± SE). All models included in the analysis are outlined in Table S1.

#### Survival model

We used a linear mixed effect model (LMM) to assess whether parental *vgll3* genotype and parental identities influenced egg and alevin survival from fertilization until the first feeding stage. The model included maternal and paternal *vgll3* genotypes and their interaction as fixed variables. Additionally, we included dam and sire IDs as random intercepts to assess the overall maternal and paternal effects on alevin survival. Interactions between dam and sire IDs were excluded from the random structure due to model convergence issues. Family location in the incubator was not included in the model as there was no temperature difference between the two incubators (Figure S1). Survival was mean-centred and standard deviation-scaled prior to fitting the model. Crosses from two females where all three or most families had a 0% survival (see Figure S5 females D25 and D33 in factorial crosses 1 and 2) were excluded from the model (6 families).

#### Female and egg trait models

We fit linear models to assess the effect of female phenotype (body length, body condition) and *vgll3* genotype on fecundity and egg traits (egg dry weight, egg lipid content, and egg protein content). All continuous variables were mean-centred and standard deviation-scaled prior to fitting the models. It should be noted that body size and body condition measured multiple months prior to spawning would give a more accurate estimate of their influence on fecundity in fishes (Koops et al., 2004). Here, we used body length and body condition at egg stripping as measures of body size and energetic condition as it was the only timepoint where this phenotypic information was available.

We used egg dry weight as a measure of egg size throughout this study as the egg wet weight measurements may be biased due to differing amount of ovarian fluid at the surface of the eggs. The egg dry weight model included the continuous variables female body length and body condition as well as *vgll3* (EE, EL, LL) genotype coded as a three-level factor variable. The egg protein model included female body length and condition and *vgll3* as fixed predictors. As we measured egg protein content as a concentration, egg dry weight was not included in the egg protein model. In addition to the same fixed predictors as the egg dry weight model, we included egg dry weight in the egg lipid model as we expected egg dry weight to influence the total amount of lipids contained in an egg.

#### Alevin trait models

We used multiple LMMs and a generalized linear mixed effect model (GLMM) to investigate associations between our traits of interest (maternal phenotype, parental *vgll3* genotypes, egg traits) and the various alevin traits. Similar to the fecundity and egg trait linear models, all continuous predictors were mean-centred and standard deviation-scaled such that each trait would have a mean of zero and a standard deviation of one before including them in the models.

Our analysis of alevin traits included body length and yolk sac area at hatching, growth rate, yolk sac consumption rate (YCR), and yolk sac conversion efficiency (YCE). All alevin trait models included maternal body length and body condition (maternal phenotype) as fixed predictors. To assess the effect of parental *vgll3* genotypes on alevin traits, we included maternal and paternal *vgll3* genotypes and their interaction as fixed factors in the models. The models also included egg dry weight and egg lipid and protein contents as fixed predictors to investigate how egg dry weight and energy content influence our alevin traits of interest.

All alevin trait models included dam ID, sire ID, and their interaction as random intercepts to partition variance into their respective maternal, paternal, and family components. In addition, the well plate location in the climate chamber was included as a random intercept to account for environmental variation within and between the two climate chambers. In the interpretation of the model results, random component variances were considered statistically important if their 95% credible interval did not overlap with zero.

Alevin body length and yolk sac area at hatching were modelled using LMMs with an identity-link function. Both models included maternal phenotype, parental *vgll3* genotypes and their interaction, and egg trait variables listed above as fixed predictors. In addition to these predictors, the hatching length and yolk sac area models included hatching date, coded as dpf (mean-centred and SD-scaled), to investigate how these hatching traits might be influenced by developmental time.

Alevin growth rate (mean-centred and SD-scaled) between hatching and 81 dpf was analysed using an LMM (identity-link). Fixed predictors included in the model were our focal traits of interest listed above (maternal phenotype, parental *vgll3* genotypes, egg traits). In addition, we included alevin body length and yolk sac size at hatching as additional predictors to investigate whether initial alevin body size or yolk energy reserves influence growth.

Yolk consumption rate (YCR; mean-centred and SD-scaled) was modelled using an LMM (identity-link) with fixed predictor model structure consisting of our focal traits of interest as well as hatching length. Length at hatching was added to the model as we wanted to investigate how initial body size influences the use of the yolk sac energy resources.

Yolk sac conversion efficiency (YCE; mean-centred and SD-scaled) was analysed using a GLMM with a Student’s t-distribution to accommodate extreme outliers of the response variable. A model assuming a Gaussian distribution was also tested, but this model could not capture the variability of the response variable as well as the GLMM. In addition to maternal phenotype, parental *vgll3* genotypes, and egg traits, we added hatching length as an additional fixed predictor to assess how initial alevin body size might influence yolk conversion dynamics.

#### Modelling technicalities and assessment of model fits

All models were fit using Bayesian statistical methods. The models used four Hamiltonian Monte Carlo chains with 5,000 iterations each and the first 500 iterations discarded as warm-up. The models resulted in 18,000 posterior samples for all parameters. Priors for the intercept, parameter estimates, and random factors were set to a relatively non-informative normal distribution with a mean of zero and standard deviation of one in all the models.

Model fits were evaluated visually by inspecting trace and autocorrelation plots which showed proper mixing of the chains and no autocorrelation. R-hat values in all models were < 1.05. Model performances were also assessed using pareto k diagnostics and posterior predictive checks. No influential points were found in the models (all k < 0.7).

#### R packages used

All data analyses were done using Rstudio version 2025.05.0 running R v4.5.0 (R Core Team, 2024). Data management and visualisation was done using the *tidyverse* package v2.0.0 (Wickham et al., 2019). The *loo* package v2.8.0 (Vehtari et al., 2017, 2024; Yao et al., 2018) was used to calculate pareto k model diagnostics. All Bayesian models were run using *rstan* v2.32.7 (Stan Development Team, 2020) interfaced via the *brms* package v2.22.0 (Bürkner, 2017, 2018, 2021).

#### Ethics statement

The Atlantic salmon broodstock used in the experiment was reared in accordance with national permits for salmon broodstocks, and no separate ethical permits were required for this study. All alevins used in the study were euthanized prior to first exogenous feeding and therefore did not require ethical approval.

## Results

### Maternal phenotype

Female body length ranged from 40.6 cm to 53.6 cm with mean length of 46.5 ± 0.6 cm across *vgll3* genotypes (Table 2). No effect of *vgll3* was found on female body length (Table S2). The mean body condition of the females at spawning was 0.88 ± 0.01 with little variation between the *vgll3* genotypes (Table 2, Table S3).

**Table 2.** Mean (± SE) fecundity, body length and condition as well as egg dry weight, lipid content and protein concentration for the maternal *vgll3* genotypes (EE, EL, LL). The number of individuals per genotype is given in parentheses after the genotype name. The overall column gives the mean (± SE) for each trait regardless of *vgll3* genotype.

| Trait | EE (*n* = 13) | EL (*n* = 11) | LL (*n* = 14) | Overall (*n* = 38) |
| --- | --- | --- | --- | --- |
| **Maternal traits** |  |  |  |  |
| Fecundity | 1854 ± 203 | 2059 ± 142 | 2147 ± 207 | 2021 ± 110 |
| Body length (cm) | 45.6 ± 0.8 | 46.6 ± 1.3 | 47.3 ± 1.1 | 46.5 ± 0.6 |
| Body condition | 0.87 ± 0.02 | 0.89 ± 0.02 | 0.88 ± 0.01 | 0.88 ± 0.01 |
| **Egg traits** |  |  |  |  |
| Egg dry weight (mg) | 35.0 ± 1.4 | 33.2 ± 2.0 | 34.5 ± 1.1 | 34.3 ± 0.8 |
| Egg lipid (mg) | 6.6 ± 0.2 | 6.5 ± 0.4 | 6.8 ± 0.3 | 6.7 ± 0.2 |
| Egg protein concentration (mg/ml) | 457.8 ± 51.7 | 436.8 ± 55.9 | 410.7 ± 33.8 | 434.4 ± 26.4 |

Female fecundity ranged from 812 to 3430 eggs (mean 2021 ± 110 eggs) (Figures S2 and S3, Table 2). No effect of female *vgll3* genotype on fecundity was found, but both body length and body condition influenced fecundity (Figures S2 and S3, Table S4). Higher body condition (model estimate: 0.30 [95% CI: 0.13, 0.46]) and longer body length (model estimate: 0.89 [95% CI: 0.73, 1.05]) were associated with higher fecundity with body length having a larger effect on egg number (Figures S2 and S3, Table S4).

No effect of maternal *vgll3* genotype was found on any of the measured egg traits as the *vgll3* parameter estimate 95% CIs overlapped zero in all of the models (Figures 4 and S4, Tables S5 – S7).

Mean egg dry weight was 34.3 ± 0.8 mg (Table 2). Neither female body condition nor body length at spawning influenced egg dry weight or lipid content (95% CIs of the estimates overlapped zero; Figure 3, Tables S5 and S6), but both influenced egg protein concentration. Longer females and those with higher body condition had lower egg protein concentration (length parameter estimate: -0.41 [95% CI: -0.64, -0.19]; condition parameter estimate: -0.31 [-0.55, -0.08]; Figure 3, Table S7).

**Figure 3.**
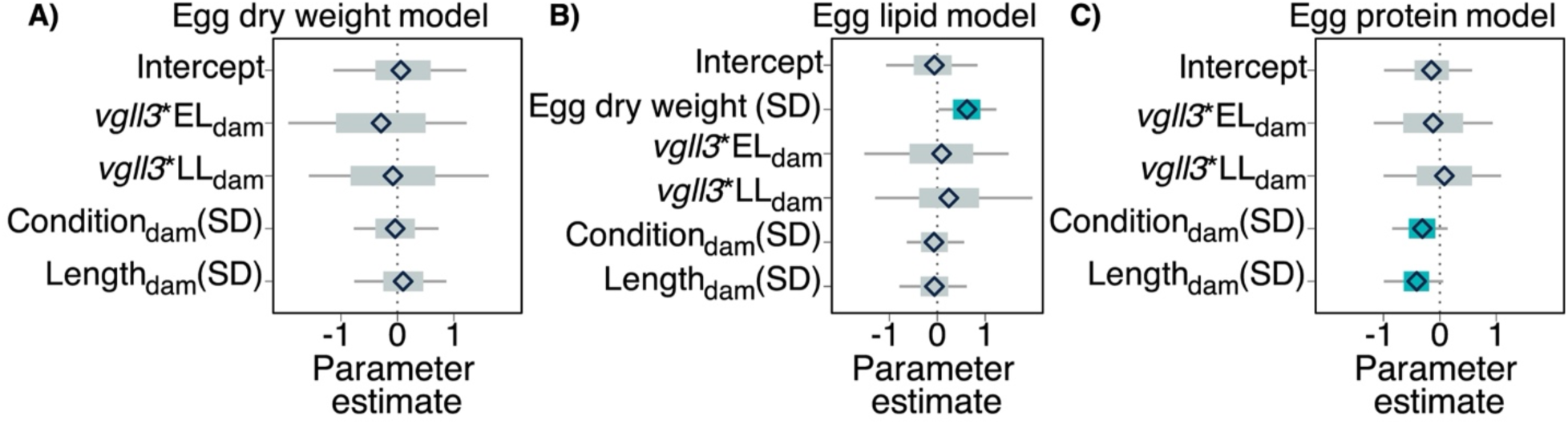
(A) Egg dry weight, (B) egg lipid and (C) egg protein model parameter estimates and their associated 95% credible intervals. For the maternal *vgll3* genotype, the contrast level is the EE genotype. Continuous were mean-centred and SD-scaled before fitting the model. The diamonds are the mean parameter estimates, the thick bars are the 95% CIs, and the thin bars are the 100% CIs. The 95% CI bars are coloured blue if the interval does not contain zero.

The mean egg lipid content across was 6.7 ± 0.2 mg (Table 2). Egg lipid content was associated with egg dry weight with heavier eggs having a higher lipid content (model estimate: 0.62 [95% CI: 0.33, 0.90], Figure 3, Table S6).

### Survival

Family-level offspring survival ranged from 0 to 89.3% (Figure S5). Embryo and alevin survival from fertilization until first feeding was largely female-dependent with higher dam compared to sire variance (Figures 4 and S5). Maternal *vgll3* genotype did not affect survival (Figure 4, Table S8). Paternal *vgll3* genotype, on the other hand, had an influence on survival where offspring of EL and LL sires had a higher survival compared to the EE sires (model estimates: 0.51 [95% CI: 0.21, 0.81] and 0.35 [95% CI: 0.06, 0.66] for the EL and LL genotypes, respectively; Figure 4, Table S8).

**Figure 4.**
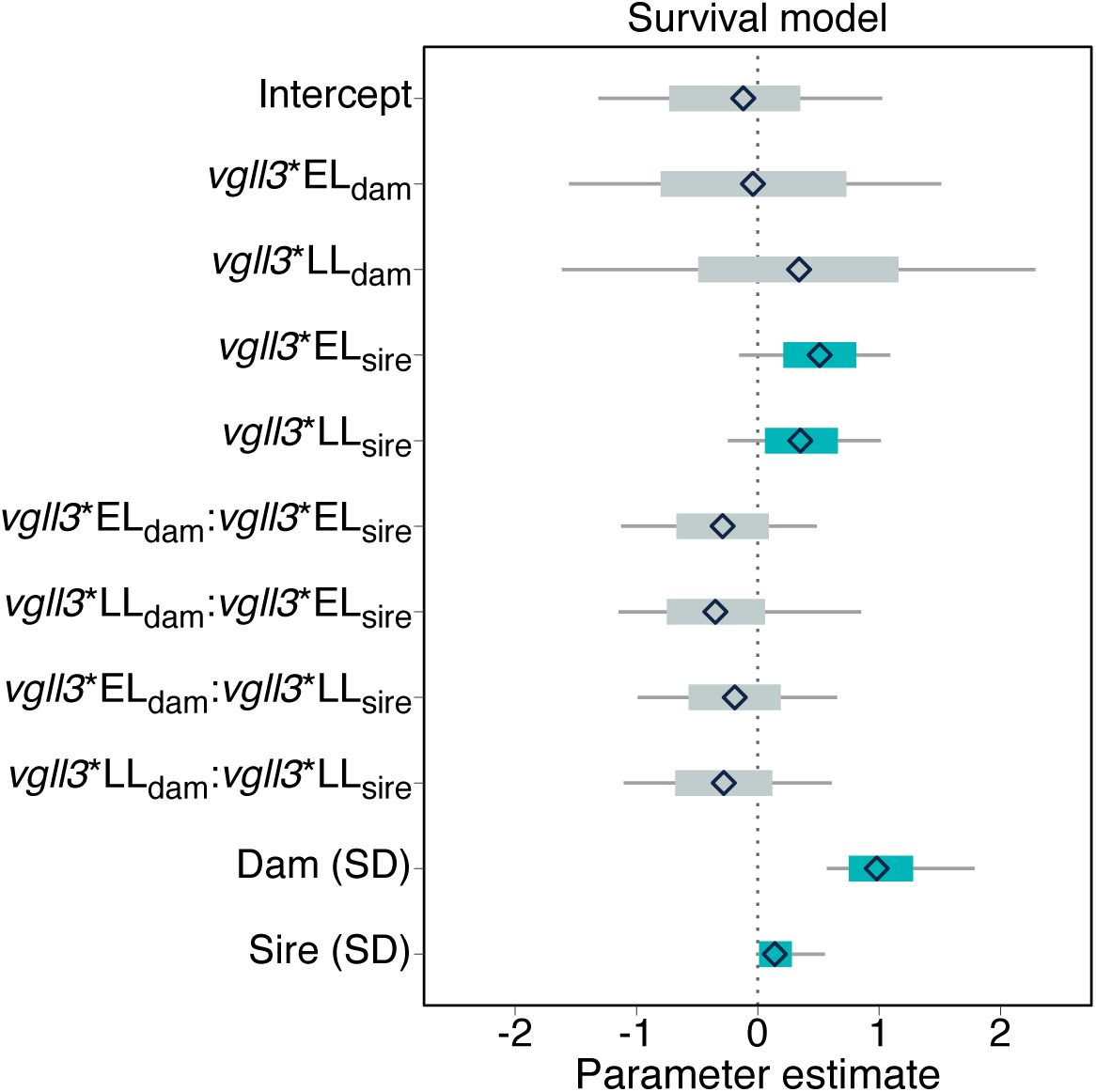
Parameter estimates and 95% credible intervals (CI) for the fixed and random effects from the alevin survival model. Survival was mean-centred and standard deviation -scaled prior to fitting the model. For the parental *vgll3* genotypes (*vgll3*_dam_, *vgll3*_sire_), the contrast level is the EE genotype. Random effect standard deviations indicate the magnitude of variation between parents (Dam and Sire). The diamonds indicate the mean parameter estimate calculated from 18,000 posterior samples, the thick bars are the 95% CIs, and the thin bars are the 100% CIs. The 95% CI bars are coloured blue if the interval does not contain zero.

### Alevin traits

Of the 960 individually reared eggs, 609 alevins hatched. Neither maternal body length nor body condition influenced any of the alevin traits that we measured in this study. In all alevin trait models, the parameter estimates for maternal body length and body condition were close to zero with 95% credible intervals that overlapped with zero (Figures S7-S10, Tables S9-S13).

Alevin body length at hatching was influenced by hatching date with alevins hatching earlier in the hatching period (56 – 68 dpf) being smaller (model estimate: 0.49 [95% CI: 0.42, 0.57]) (Figures S6 and S7, Table S9). Egg size measured as within-female average egg dry weight did not have a major effect on hatching length of individually reared alevins from the same family (the parameter estimate 95% CI overlapped with zero), but egg dry weight did influence yolk sac size at hatching (Figures 5, S7, and S8, Tables S9 and S10). The alevins that hatched from heavier eggs had larger yolk sacs (model estimate: 0.73 [95% CI: 0.34, 1.10]). Egg lipid and protein contents did not influence alevin length or yolk size at hatching with their respective parameter estimates being close to zero (Figures S7 and S8, Tables S9 and S10).

**Figure 5.**
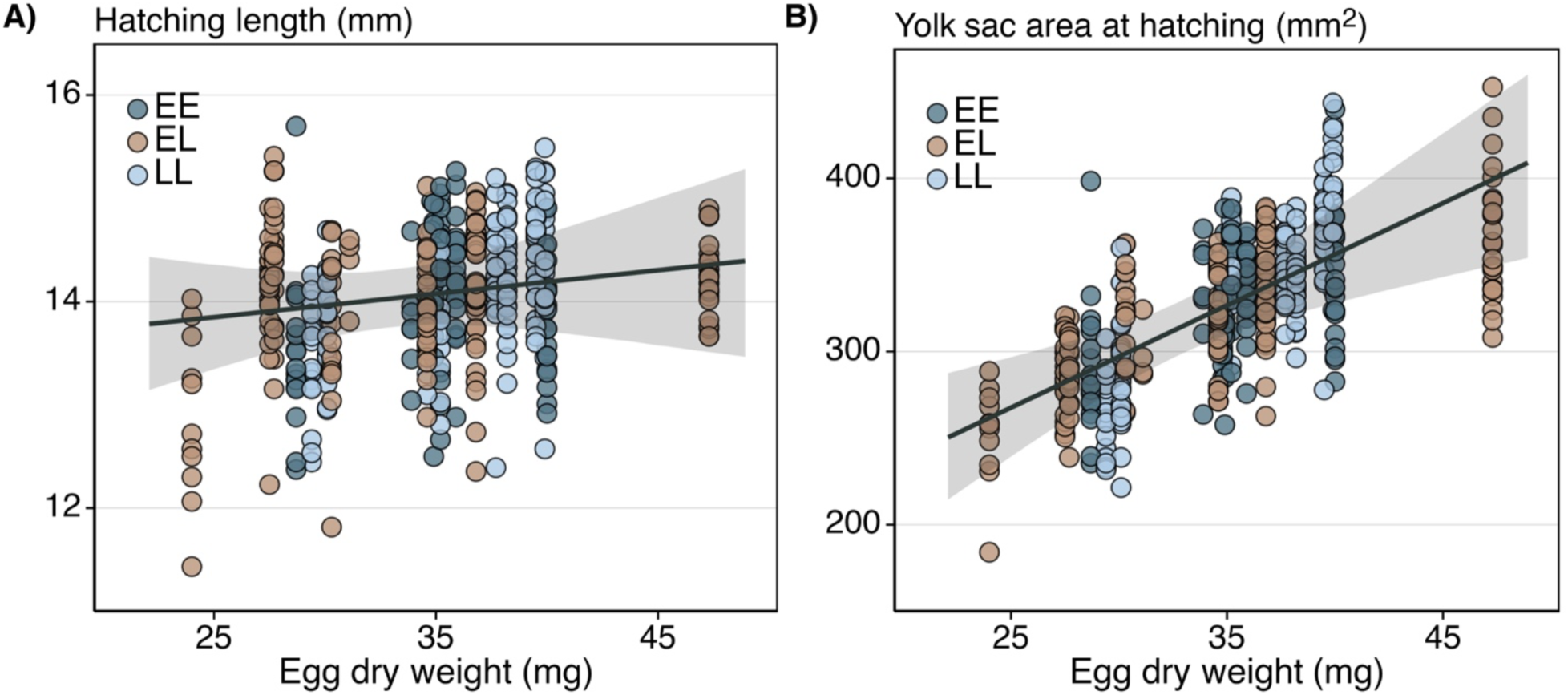
(A) Alevin body length (mm) and (B) yolk sac area (mm^2^) at hatching plotted against egg dry weight (mg). Egg dry weight is the female average egg dry weight. Each point represents a single alevin with measurement taken within 24 hours of hatching. The points are coloured according to the maternal *vgll3* genotype (dark blue = EE, light brown = EL, light blue = LL). The trendlines are the predicted hatching length and yolk sac area calculated from the respective models. For the predictions, parental *vgll3* genotypes were set to EL and maternal body length and condition, egg lipid and protein contents and hatching date were set to the mean. The shaded area around the trendline indicates the 95% credible interval.

Neither maternal nor paternal *vgll3* genotype influenced alevin hatching length or yolk sac size (Figures S7 and S8 and Tables S8 and S9).

We measured alevin growth between hatching and 81 dpf as an increase of alevin body length in millimetres per day. Hatching length influenced growth rate with alevins that hatched at a larger size having a slower growth rate compared to alevins that were smaller at hatching (model estimate: -0.37 [95% CI: -0.52, -0.23]; Figures 7 and S9, Table S11). None of our measured egg traits had a major influence on growth rate (Table S11). Growth was not influenced by parental *vgll3* genotypes and the parameter estimates for *vgll3* were close to zero with their associated 95% CIs overlapping with zero (Table S11).

Length at hatching had a slight influence on alevin yolk sac consumption rate (Figure S10) with larger alevins consuming their yolk sac energy reserves at a higher rate (model estimate: 0.13 [95% CI: 0.00, 0.27]; Table S12). Of the measured egg traits, egg lipid content affected alevin yolk consumption. Larger egg lipid content was associated with a lower yolk consumption rate (model estimate: -0.30 [95% CI: -0.59, -0.03]; Figure 6, Table S12]). No effects of parental *vgll3* genotype were found on yolk consumption rate (model parameter estimate 95% CIs overlapped zero, Figure S10, Table S12).

**Figure 6.**
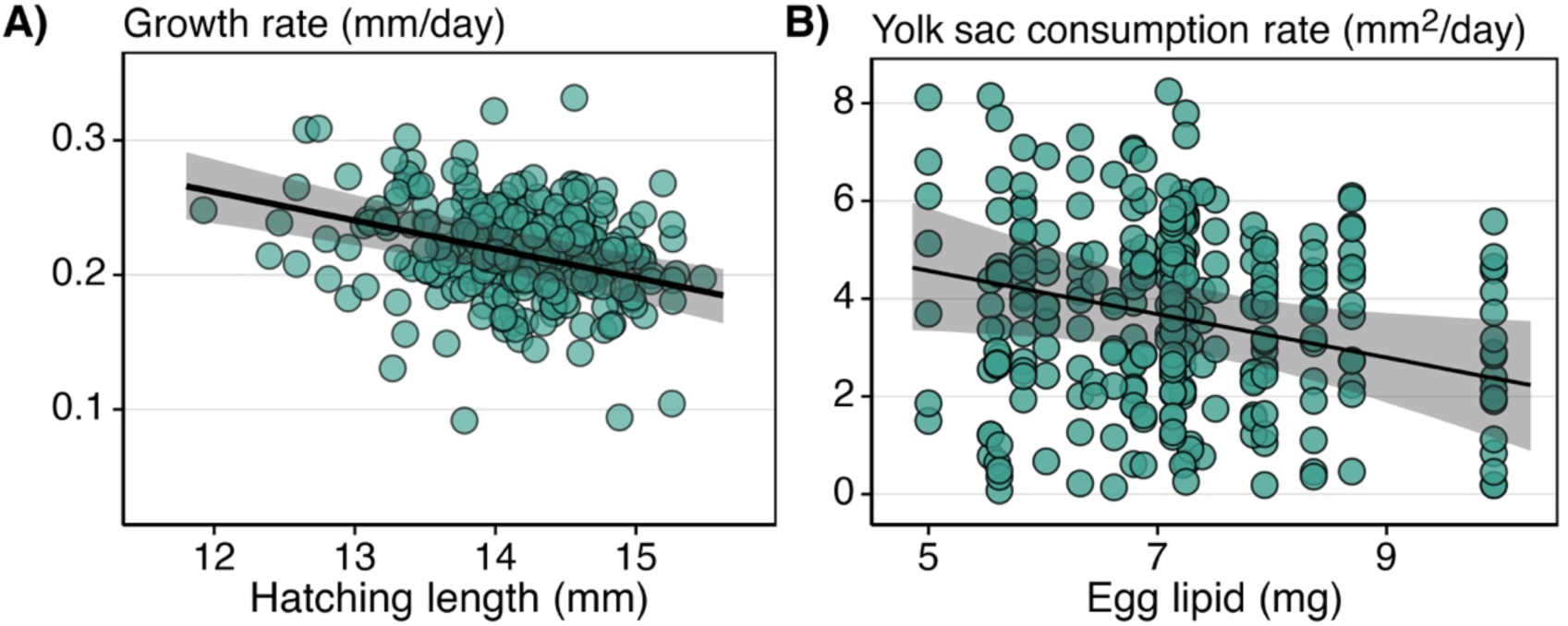
A) Alevin growth rate (mm/day) plotted against body length at hatching (in mm) and B) yolk sac consumption rate (mm^2^/day) plotted against egg lipid content (mg). The trendlines are the predicted growth rate and yolk sac consumption rate calculated from their respective models. For the growth predictions, parental *vgll3* genotypes were set to EL, maternal body length and condition, all egg traits, and alevin yolk sac area at hatching were set to the mean. For the yolk sac consumption predictions parental *vgll3* genotypes were set to EL and maternal body length and condition, egg lipid and protein contents and hatching length were set to the mean. The shaded areas around the trendlines indicates the 95% credible interval.

None of our main factors of interest (i.e., maternal phenotype, parental *vgll3* genotypes or egg traits) influenced yolk conversion efficiency (Table S10) with the majority of the alevins having a very similar conversion efficiency (Figure S11).

### Parental effects on alevins in their early life development

We assessed maternal and paternal effects on alevin traits in all models as the among-female and among-male variation obtained from the model random factors. Paternal effects on all measured alevin traits were negligible with the estimate-associated 95% CIs bounded at zero or close to zero (Figure 7, Tables S9 - S13). Maternal effects, on the other hand, were present for most alevin traits. The among-female variation for length and yolk sac area at hatching were 0.51 [95% CI: 0.31, 0.79] and 0.43 [95% CI: 0.27, 0.66], respectively. For alevin growth and yolk consumption rate, the variation among females was 0.34 [95% CI: 0.06, 0.63] and 0.21 [95% CI: 0.01, 0.47], respectively. Variation among females for yolk conversion efficiency was mostly absent (dam variance: 0.02 [95% CI: 0.00, 0.05]). Similar to paternal effects, among-family variation, estimated as an interaction between maternal and paternal identities, was negligible with the estimate 95% CIs bounded at zero or very close to zero (Figure 7, Tables S9 - S13).

**Figure 7.**
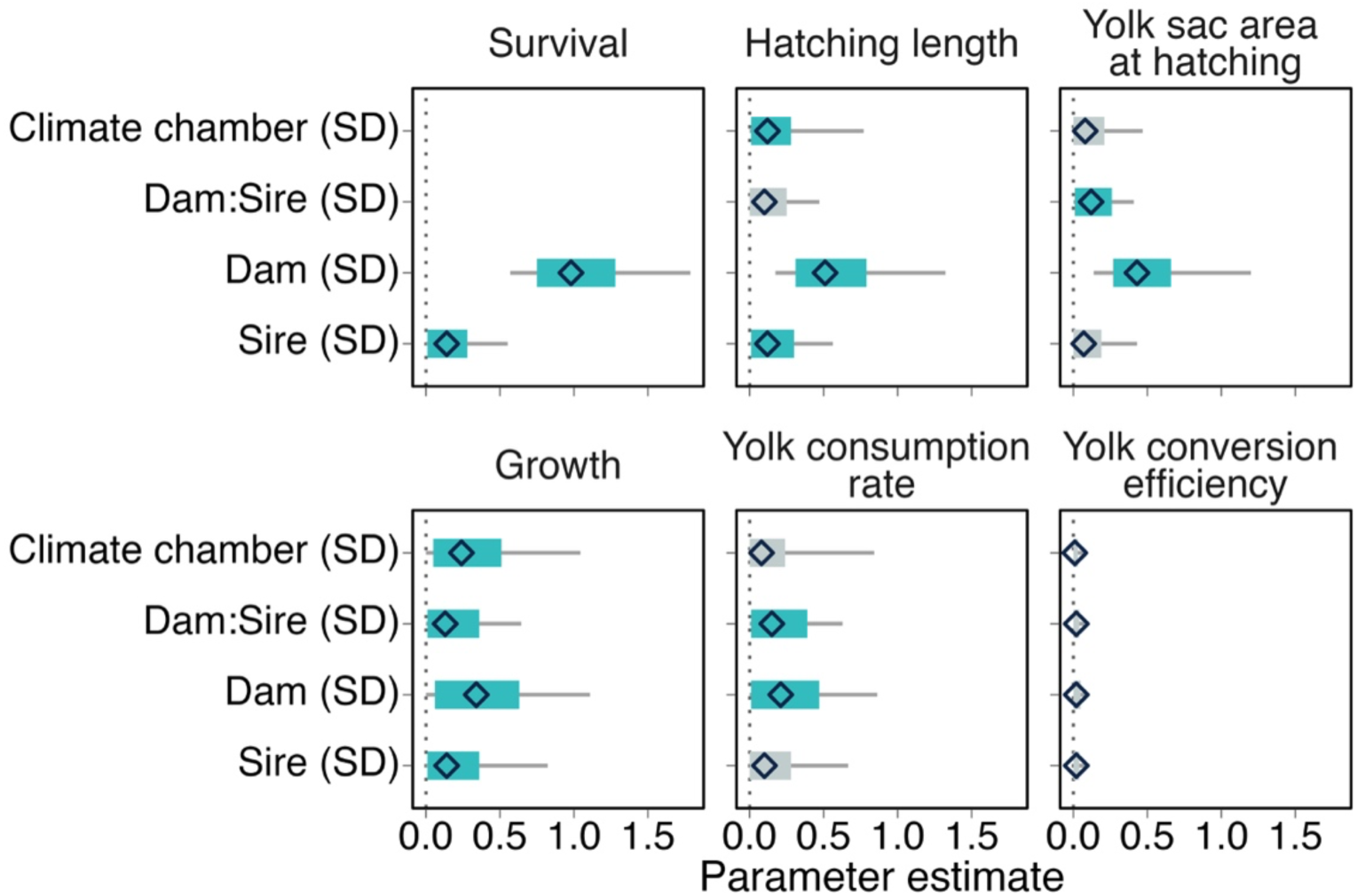
Random effect parameter estimates from alevin survival, hatching length and yolk sac area, growth, yolk consumption rate and yolk conversion efficiency models. Random effect standard deviations indicate the magnitude of variation between climate chambers and families (Dam:Sire) and between parents (Dam and Sire). The diamonds indicate the mean parameter estimate calculated from 18,000 posterior samples, the thick bars are the 95% CIs, and the thin bars are the 100% CIs. The 95% CI bars are coloured blue if the interval does not contain zero.

## Discussion

In this study we explored the influences of parental genotypes of a maturation-related gene, maternal phenotype, and egg traits on various offspring traits to understand how parents, and mothers in particular, influence offspring survival, growth, and development. By performing crosses based on the parental *vgll3* genotypes, we were able to disentangle the maternal and paternal contributions to variation in offspring early life. Furthermore, we measured multiple egg traits to understand how maternal egg energy provisioning influences offspring phenotypic traits. We did not find an effect of the *vgll3* genotype of either parent on the majority of our measured alevin phenotypic traits. The only association between parental *vgll3* genotype and the offspring was a genetic effect of paternal *vgll3* genotype on offspring survival. Egg traits affected alevin hatching phenotype and yolk sac usage highlighting the importance of maternal provisioning in offspring early life. We demonstrate that offspring early-life traits were largely influenced by a variety of maternal effects unrelated to maternal *vgll3* genotype, whereas the paternal contribution was limited.

### Maternal phenotype influences egg energy provisioning, but not size

Larger females, in general, produce larger eggs in various fish species (Barneche et al., 2018; Hixon et al., 2014; Koenigbauer & Höök, 2023), though the strength of this relationship can vary (Heinimaa & Heinimaa, 2004; Maamela et al., 2023). We did not find an effect of female body length or body condition on egg size as measured by egg dry weight. This could be due to all of our females sharing a common rearing environment in the hatchery and being the same age and thus, being relatively similar in size at spawning. A wider range of phenotypes (e.g., size, age, number of spawning events) is likely needed to better disentangle female body size effects on egg size.

Female body size and body condition can also influence egg energy content with larger and better conditioned females provisioning their eggs with more energy (Heinimaa & Heinimaa, 2004; Ouellet, 2001). In contrast, we found an opposite effect of female body size and condition on egg energy content when measured by egg protein concentration. Larger females and those with higher body condition had lower egg protein concentration. Female body size or condition did not affect egg lipid content, but larger eggs did hold more lipids. These female body size and condition results indicate that the females were allocating energy to somatic growth rather than egg energy provisioning.

Female *vgll3* genotype had no influence on any of the egg traits we measured. Our expectation was that if maternal *vgll3* genotype affects egg traits, it would have likely been via egg lipid content. Previously, *vgll3* has been found to influence body condition (Debes et al., 2021; House et al., 2023) and lipid traits (House et al., 2025) in juvenile Atlantic salmon. Body condition is related to body lipid reserves and thus, is a proxy for available energy reserves in salmon that the female can use to provision eggs (Shearer & Swanson, 2000). The lack of an effect of *vgll3* genotype on egg lipid content indicates that the allocation of lipids to eggs is unlikely mediated via *vgll3* genotype, at least among hatchery-reared females.

### Parents influence egg and alevin survival

We observed high variation in egg and alevin survival up to first feeding with family-level survival ranging from 0% up to 89%. Survival was largely influenced by maternal effects likely linked with egg quality. Egg size and nutritional composition can affect both embryo and larval survival in fish (Brooks et al., 1997; Kaththriarachchi et al., 2025; Lehnert et al., 2018; Srivastava & Brown, 1991; Tveiten et al., 2004; Wootton & Smith, 2015). We measured survival only between fertilization and first feeding and therefore, cannot determine what the precise cause of mortality was for fertilized eggs. It is unlikely that fertilization success explains the variation in survival estimates to a large extent as we removed visibly unfertilized eggs from the incubators the day after fertilizations and these removed eggs were not included in the survival estimates. Despite removing the unfertilized eggs, the cause for complete failure in survival in some families could still be an artefact of the fertilization process rather than complications during post-fertilization development as none or very few of the eggs developed. Potential causes of egg and alevin mortality could be related to complications in development leading to malformations during the embryonic and alevin stages, such as the alevin not being able to utilize its yolk resources (Bobe & Labbé, 2010; Bonnet et al., 2007; Eriksen et al., 2006). At first feeding, alevins still had a small amount of yolk left, which indicates that starvation likely wasn’t a major cause of mortality.

It should be noted that egg and alevin survival and its possible link with egg quality might not be solely related to a female’s inherent egg characteristics or quality, but rather it may also relate to the timing of egg stripping and post-ovulatory ageing. Female Atlantic salmon have group synchronous ooycyte development (Wootton & Smith, 2015). In captivity, Atlantic salmon do not spawn without access to suitable spawning habitat and mature salmon need to be manually stripped (Fleming & Einum, 2011). Delaying egg stripping post-ovulation can increase egg and alevin mortality due to egg overripening (Aegerter & Jalabert, 2004; de Gaudemar & Beall, 1998; Mommens et al., 2015). It is therefore possible that some of the females with very low family-level survival over all crosses with males could have had overripe eggs.

In our study, paternal, but not maternal, *vgll3* genotype influenced survival up to first feeding with offspring of EL or LL sires having higher survival compared to offspring of EE sires. As we observed the gene’s effect on survival during the egg and endogenous feeding period, the results suggest that paternal *vgll3* genotype influences alevin physiology before behavioural traits such as resource acquisition emerge. It is unclear how paternal *vgll3* genotype may influence offspring survival. However, Aykanat et al., (2024) found that parental *vgll3* genotypes can affect early-life survival of post-first-feeding Atlantic salmon in the wild. Their study found that parental L alleles were associated with higher survival although the study did not explicitly differentiate whether this *vgll3* effect was mediated more by the maternal or paternal *vgll3* genotype. Nevertheless, the results of the study by Aykanat et al., (2024) and our study indicate that parental, and more specifically paternal, *vgll3* genotypes can influence offspring early survival both prior to and during independent feeding. However, in our experiment, we did not take into consideration the *vgll3* genotype of the offspring as individual genotype data is not available for the alevins used in the analysis of survival up to first feeding. Estimating the contributions of both parental and offspring *vgll3* genotypes would help to disentangle the mechanisms behind this gene’s potential influence on offspring survival.

### Egg traits and hatching phenotype are associated with offspring early development

We found a positive effect of egg size on alevin body size at hatching. This result is in line with previous studies that found that larger eggs give rise to larger offspring in a wide range of species (Cogliati et al., 2018; Einum & Fleming, 1999; Fischer et al., 2002; Leblanc et al., 2023; Martin & Pfennig, 2010; Paulsen et al., 2009; Wu et al., 2022). Interestingly, egg size only affected yolk sac size at hatching and not body length where alevins that hatched from larger eggs had larger yolk sacs. Thus, our results indicate that the potential benefit of larger egg size to alevins (Einum & Fleming, 2000) is likely in the form of greater available resources overall in early life. This egg size benefit is not tied to egg lipid or protein content (female averages) specifically, given that these traits were not associated with alevin size at hatching.

Hatching day influenced alevin body length at hatching. Earlier hatching alevins were shorter compared to later hatching alevins indicating a faster embryonic developmental rate of the smaller hatching individuals. Despite the finding that hatch date influences body size, there was little variation in hatching day overall in our study and the majority of the alevins hatched within a short period of time (∼3 days). Similar findings of small variation in hatching time have been reported in salmonids, when eggs are incubated at similar temperatures (Leblanc et al., 2016; Van Leeuwen et al., 2016) suggesting that hatching time is largely dependent on temperature regime but that smaller individuals tend to hatch earlier.

In fish, early-life growth rate can be influenced by egg size, the time of hatching, and initial body size at hatching, which may all be influenced by environmental conditions (Gilbey et al., 2009; Heath et al., 1999; Penney et al., 2022; Self et al., 2018; Simonin et al., 2016). For example, smaller offspring or offspring hatching from smaller eggs can grow faster than larger offspring (Gilbey et al., 2009; Heath et al., 1999; Penney et al., 2022; Self et al., 2018; Simonin et al., 2016). We found a similar association between initial body length and growth rate where higher growth rate was observed in alevins that hatched at a smaller size. The higher growth rate of smaller offspring may be linked to compensatory growth due to limited energy resources (Ali et al., 2003; Metcalfe & Monaghan, 2001). If the higher growth rate of smaller alevins was connected to limited energy resources, we would expect to see an association between initial yolk sac size and growth. We did not find such an effect of yolk sac size indicating that compensatory growth is probably not driving the higher growth rate in our study.

Another explanation of the observation that smaller alevins have higher growth rate may be related to alevin fitness. Larger offspring body size is generally thought to result in higher fitness (Kingsolver & Pfennig, 2004; Mobley et al., 2020). In salmon, the timing of independent feeding when juveniles emerge from redds after depleting their yolk sacs is one of the major sources of mortality in salmon early life (Fleming & Einum, 2011). Larger size at emergence can lead to higher survival due to, for example, competitive advantage and larger somatic energy stores (Einum & Fleming, 1999, 2000; Hutchings, 1991; Johnsson et al., 1999). With this fitness advantage in mind, the higher growth rate of smaller hatching alevins could be an indication of alevins maximizing their growth and subsequent body size from early on in life to be better equipped for the exogeneous feeding period. We were not able to follow individual-level alevin growth up to independent feeding. Therefore, we do not know whether the smaller hatching alevins would have been able to maintain their higher growth throughout the alevin stage and how initial size at hatching would have affected size at independent feeding and beyond.

Beyond understanding how initial alevin body and yolk sac size influence growth, we were interested in how egg size and energy composition influence growth and yolk sac use dynamics (Thorn et al., 2019). None of our measured egg traits influenced alevin growth. We did, however, observe an effect of egg lipid content on yolk consumption. Alevins whose mother produced eggs with higher lipid content consumed their yolks at a slower rate. The energy density of lipids is higher compared to protein (Kamler, 2008). Therefore, less of the yolk energy resources are needed for basic maintenance metabolism and higher egg lipid content and lower yolk consumption rate could buffer alevins against starvation-induced mortality (Einum, 2003). Maternal diet can influence egg energy composition (Maamela et al., 2023; Rennie et al., 2005) which emphasizes the link between maternal resource acquisition opportunities and alevin early life development.

Alevin yolk sac conversion efficiency was not influenced by any of our measured parental, egg or alevin traits. Similarly, previous studies were unable to find clear differences in yolk conversion efficiency in wild, domesticated or hybrid Atlantic salmon (Debes et al., 2013; Solberg et al., 2014). Our results, together with the previous findings, indicate a lack of overall variability and initial energy resource effect on yolk sac conversion efficiency in stable conditions.

### Maternal effects are prominent in offspring early life

By estimating the contributions of the males and females in our models, we could estimate the strength of maternal and paternal effects on offspring survival and developmental traits. Apart from the paternal *vgll3* genotype influencing survival, paternal effects were largely absent for all our traits of interest. Prior evidence for paternal effects in salmonids is limited or lacking (Einum, 2003; Nadeau et al., 2009; Páez et al., 2010; Yamamoto et al., 2021), though there are exceptions (M. L. Evans et al., 2019; Houde et al., 2011; Pakkasmaa et al., 2001). Our results suggest that paternal effects are unlikely to contribute significantly to variation in survival, growth, and early development of Atlantic salmon in stable conditions. On the other hand, we did find an influence of the mother on most of the measured alevin traits. Importantly, these maternal effects or unidentified attributes of the mother were found to influence offspring traits in addition to the effects of maternal body length, body condition, and egg size and nutritional content. Maternal effects were greater for traits associated with hatching (e.g., length at hatch) and survival, and smaller for traits related to yolk sac usage (e.g., yolk consumption rate). Offspring traits expressed post-hatching such as growth and use of the yolk resources are dependent on not only the maternally provided resources, but they are also potentially influenced by the offspring’s own genotype and physiology. Taken together, the relatively small maternal effects associated with the alevin developmental traits support the notion that maternal effects decrease during ontogeny in a variety of species including salmonids (Gilbey et al., 2009; Houde et al., 2013, 2015; Moore et al., 2019; Perry et al., 2004; Van Leeuwen et al., 2016) and that the strength of maternal (and paternal) effects is timepoint-specific.

### Conclusions

Overall, we show that in a species with minimal parental care, mothers have a great capacity to influence their offspring early life development via maternal provisioning of eggs. Although we did not find strong evidence of paternal effects to affect our measured offspring traits, the finding of paternal genetic effect on offspring survival illustrates that fathers may play an important role in determining offspring fitness. Other than this paternal *vgll3* genetic effect on survival, parental *vgll3* genotype effects were minimal on the offspring traits measured here, but future research incorporating estimates of parental versus offspring genetic contributions to offspring fitness would further disentangle the role parents play in shaping their offspring’s life.

## Author contributions

Conceptualization: KSM, CRP, KBM; Data curation: KSM; Formal analysis: KSM; Funding acquisition: KSM, CRP; Investigation: KSM, JMP, CS, XDH, CRP; Methodology: KSM, JMP, CS, XDH, CRP, KBM; Project administration: CRP; Resources: JMP, CRP; Supervision: CRP, KBM; Visualization: KSM; Writing-original draft: KSM; Writing-review & editing: KSM, JMP, CS, XDH, CRP, KBM

## Acknowledgements

We thank Dr. Lucy Cotgrove, Tiia Leinonen, and Luke Taivalkoski hatchery staff for help with gamete collection, Dr. Amaïa Lamarins and Gautier Magne for help with performing the crosses, Nikolai Piavchenko for help with animal husbandry, and Dr. Iikki Donner and Titta Liukkonen for help with sampling during the experiment. Dr. Frédéric Guillaume is thanked for providing access to the climate chambers used to rear the alevins, Juhani Maamela for providing the photographing equipment, Dr. George Hancock for creating the ImageJ macro used to measure alevin phenotypic traits, and Dr. Andrew House for giving advice and guidance during the lipid extractions.

## Data availability statement

All data and code used in the analysis will be made available in Zenodo via the URL https://doi.org/10.5281/zenodo.18620194 upon acceptance of the article.

## Supplementary material for

**Figure S1.**
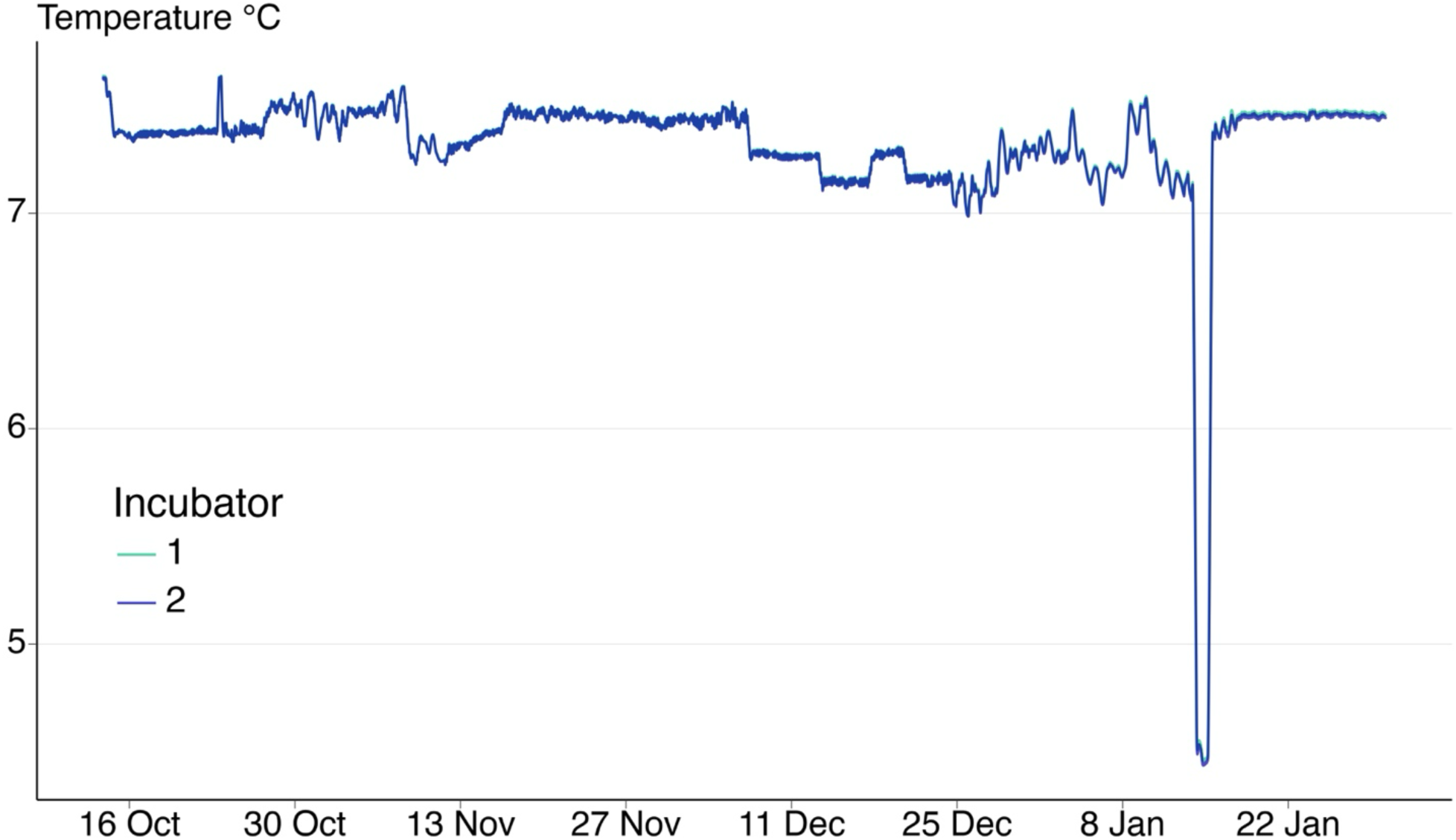
Water temperature (°C) in the incubators from fertilization until first-feeding stage. The two lines indicate the two day rolling average temperature in the two incubators (green = incubator 1, purple = incubator 2). The temperatures in the two incubators did not differ during the incubation and alevin period (note the green line behind the purple line). On 14 January 2024 (93 dpf), there was a water temperature control system failure in the incubators, which caused a drop in water temperature down to approximately 4.5°C for one day.

**Figure S2.**
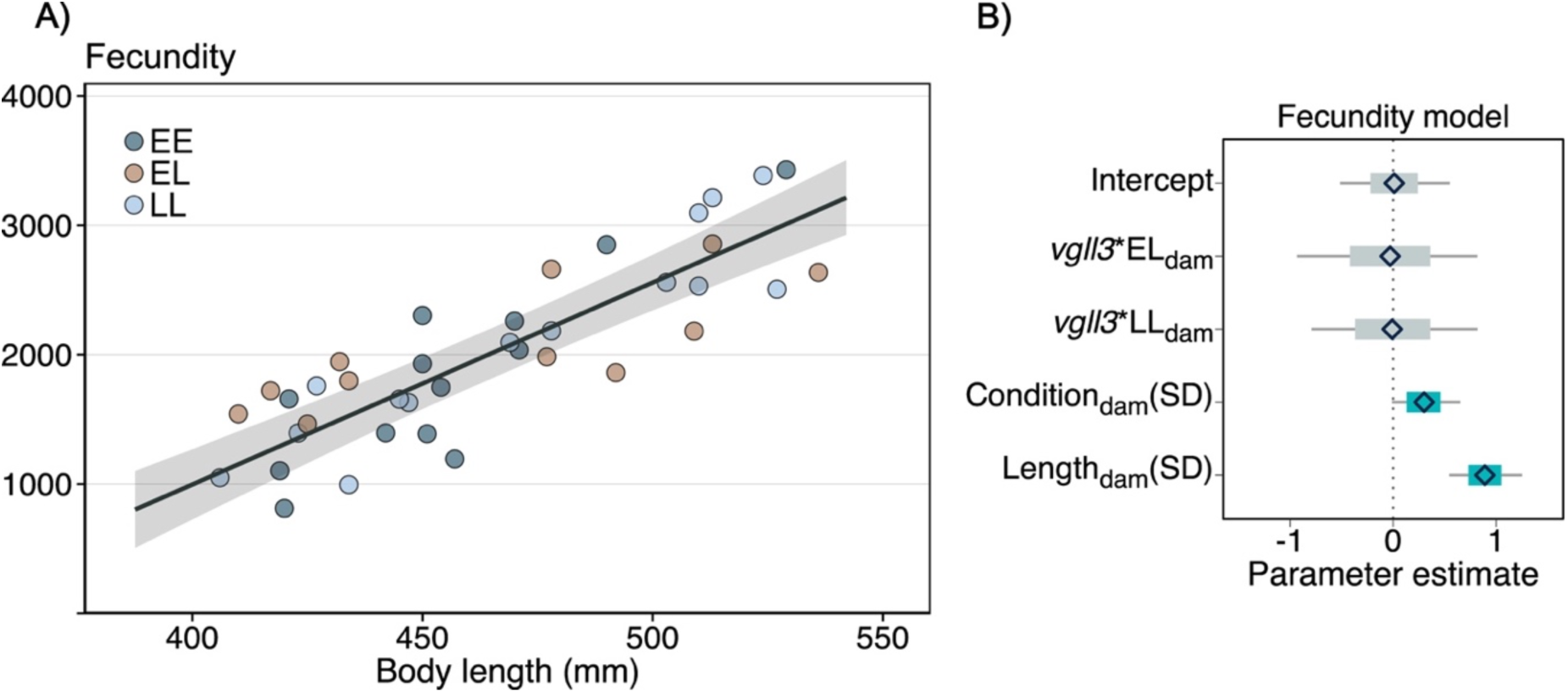
(A) Fecundity of mature female Atlantic salmon plotted against body length at spawning. Points indicate observed fecundities coloured according to the *vgll3* genotype of the female (dark blue = EE, light brown = EL, light blue = LL). The trendline is the predicted fecundity calculated from the fecundity model. For the predictions, *vgll3* genotype was set to EL and body condition to the mean body condition of the females. The shaded area around the trendline indicates the 95% credible interval. (B) Fecundity model mean parameter estimates and their associated 95% credible intervals. For the maternal *vgll3* genotype, the contrast level is the EE genotype. Body condition and length were mean-centred and SD-scaled before fitting the model. The diamonds are the mean parameter estimates, the thick bars are the 95% CIs, and the thin bars are the 100% CIs. The 95% CI bars are coloured blue if the interval does not contain zero.

**Figure S3.**
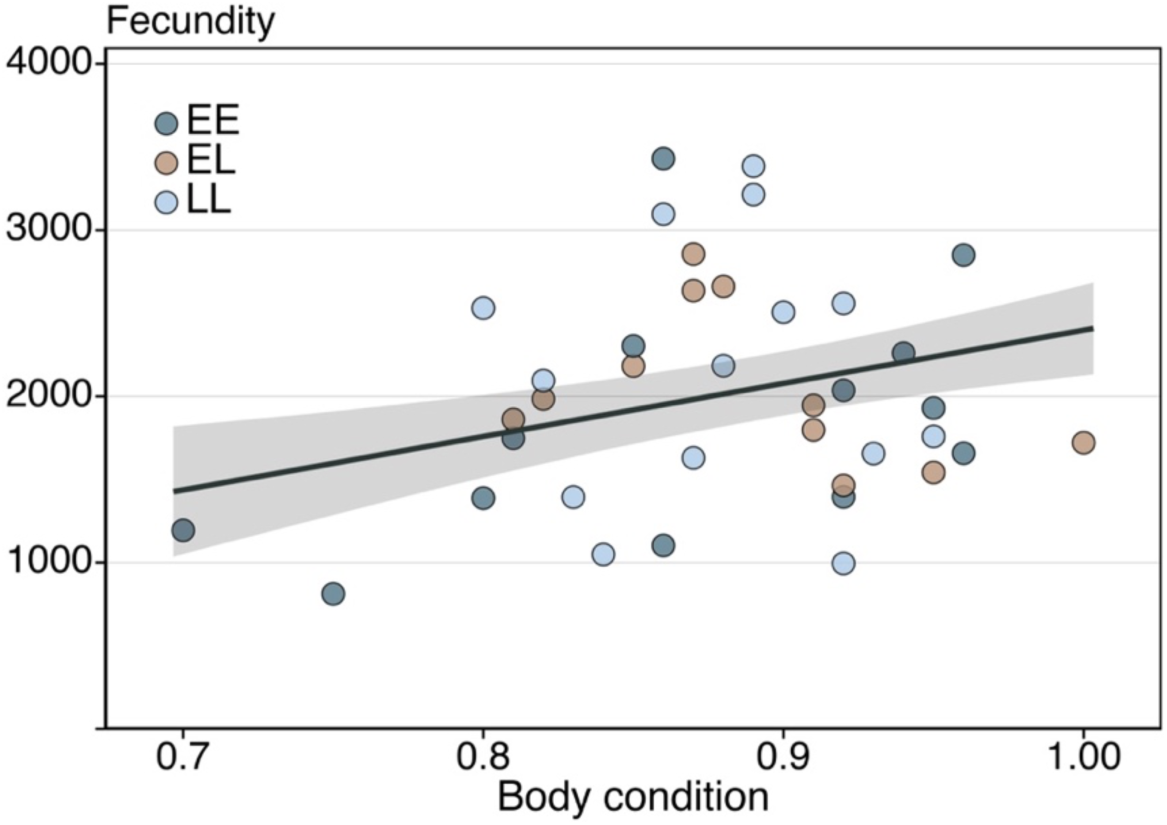
Fecundity of mature female Atlantic salmon plotted against body condition at spawning. Points indicate observed fecundities coloured according to the *vgll3* genotype of the female (dark blue = EE, light brown = EL, light blue = LL). The trendline is the predicted fecundity calculated from the fecundity model. For the predictions, *vgll3* genotype was set to EL and body length to the mean length of the females included in the model. The shaded area around the trendline indicates the 95% credible interval.

**Figure S4.**
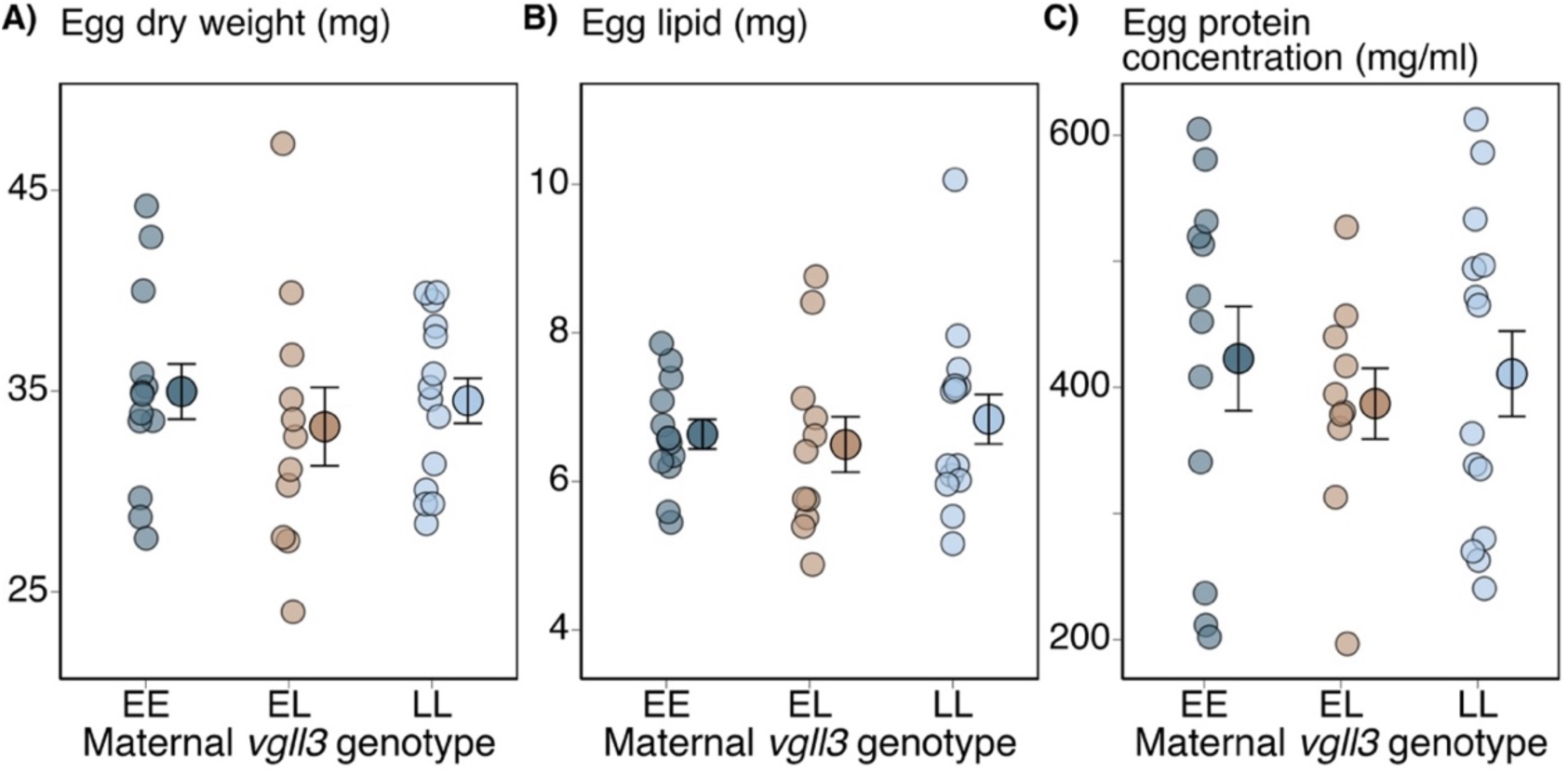
(A) Egg dry weight, (B) egg lipid content, and (C) egg protein concentration plotted against maternal *vgll3* genotype. The smaller points are the observed data for individual females, whereas the larger points with the error bars indicate the mean and standard error of the mean.

**Figure S5.**
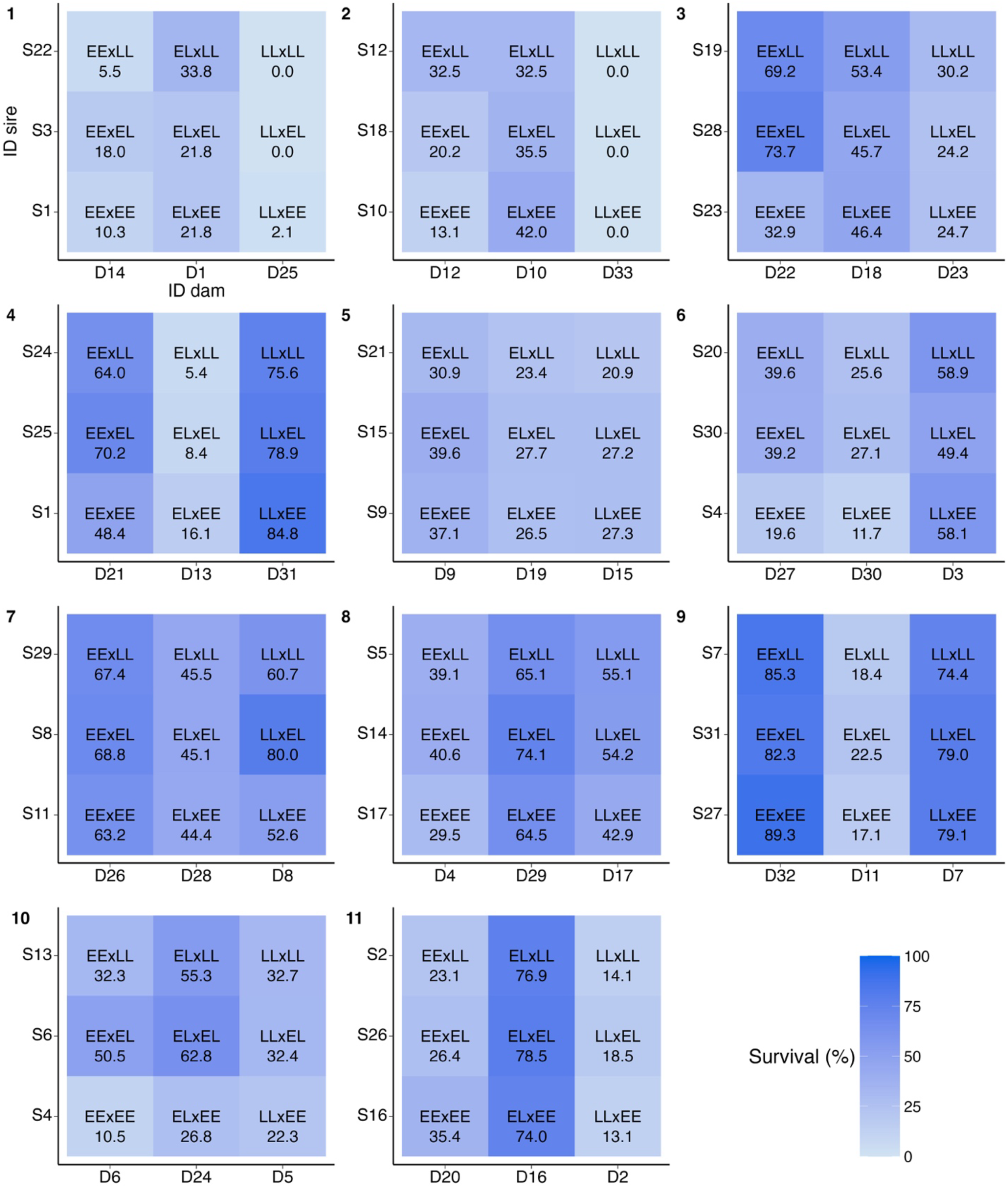
Family-level alevin survival (%) in the incubators. Each square represents a unique 3 x 3 factorial cross based on the parental *vgll3* genotypes. Within a square, individual dams are in the columns and individual sires are on the rows. The genotypes inside the coloured squares tell the parental *vgll3* genotypes of that specific cross with the maternal *vgll3* genotype listed first. The value below the parental genotypes is the percent survivorship from fertilization (0 dpf) until first feeding (110 dpf).

**Figure S6.**
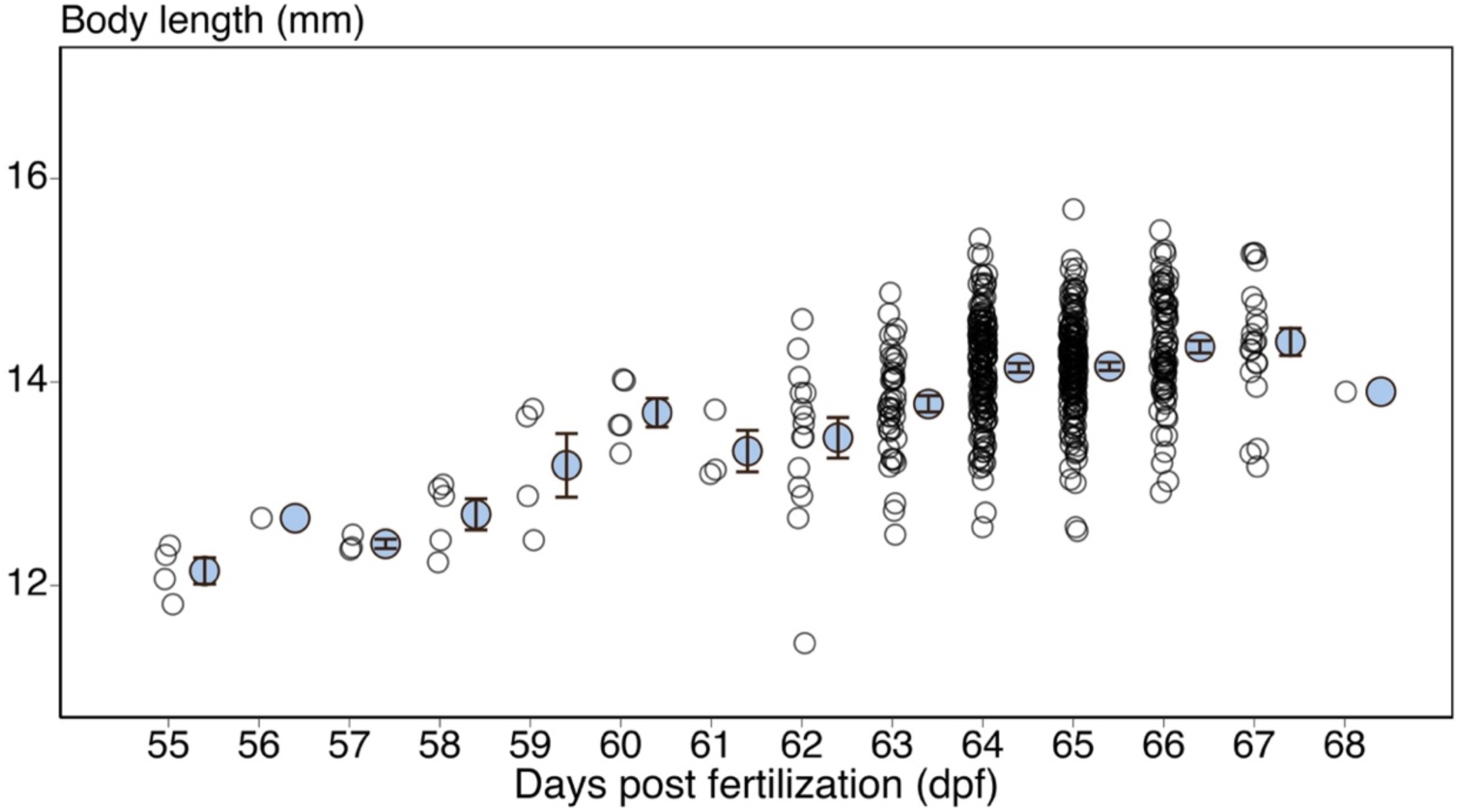
Alevin body length (mm) at hatching plotted against hatching day (dpf). Open circles denote body length measurement from an individual alevin taken within 24 hours of hatching. The blue circles indicate the average alevin body length within a single day and the error bars indicate SE. Peak hatching of alevins occurred at 64 – 66 dpf.

**Figure S7.**
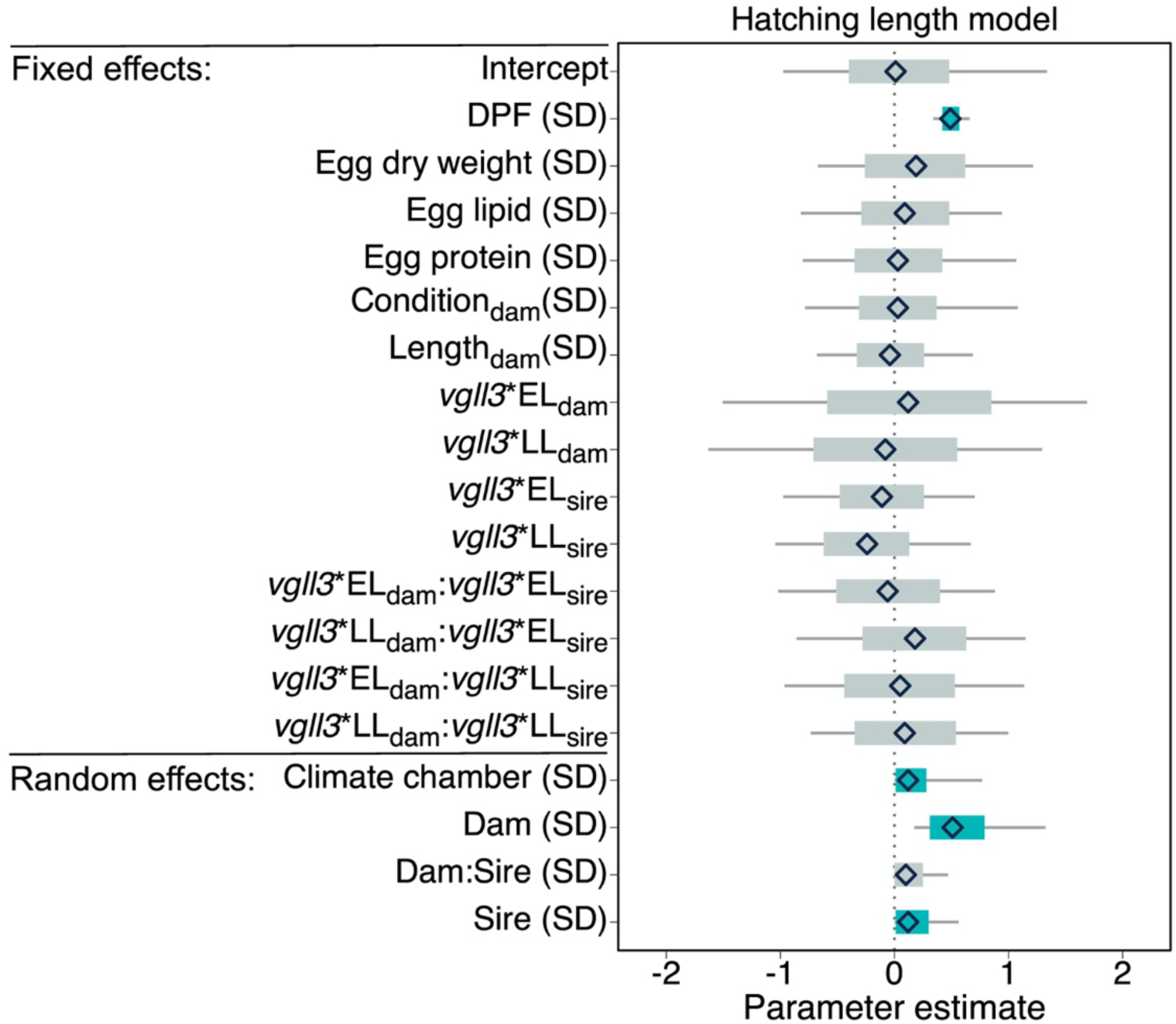
Parameter estimates and 95% credible intervals (CI) for the fixed and random effects from the alevin hatching length model. All continuous variables were mean-centred and standard deviation -scaled prior to fitting the model. Thus, the model estimates indicate the effect of increasing or decreasing the variable by one SD. For the parental *vgll3* genotypes (*vgll3*_dam_, *vgll3*_sire_), the contrast level is the EE genotype. Random effect standard deviations indicate the magnitude of variation between climate chambers and between parents (Dam and Sire) and families (Dam:Sire). The diamonds indicate the mean parameter estimate calculated from 18,000 posterior samples, the thick bars are the 95% CIs, and the thin bars are the 100% CIs. The 95% CI bars are coloured blue if the interval does not contain zero.

**Figure S8.**
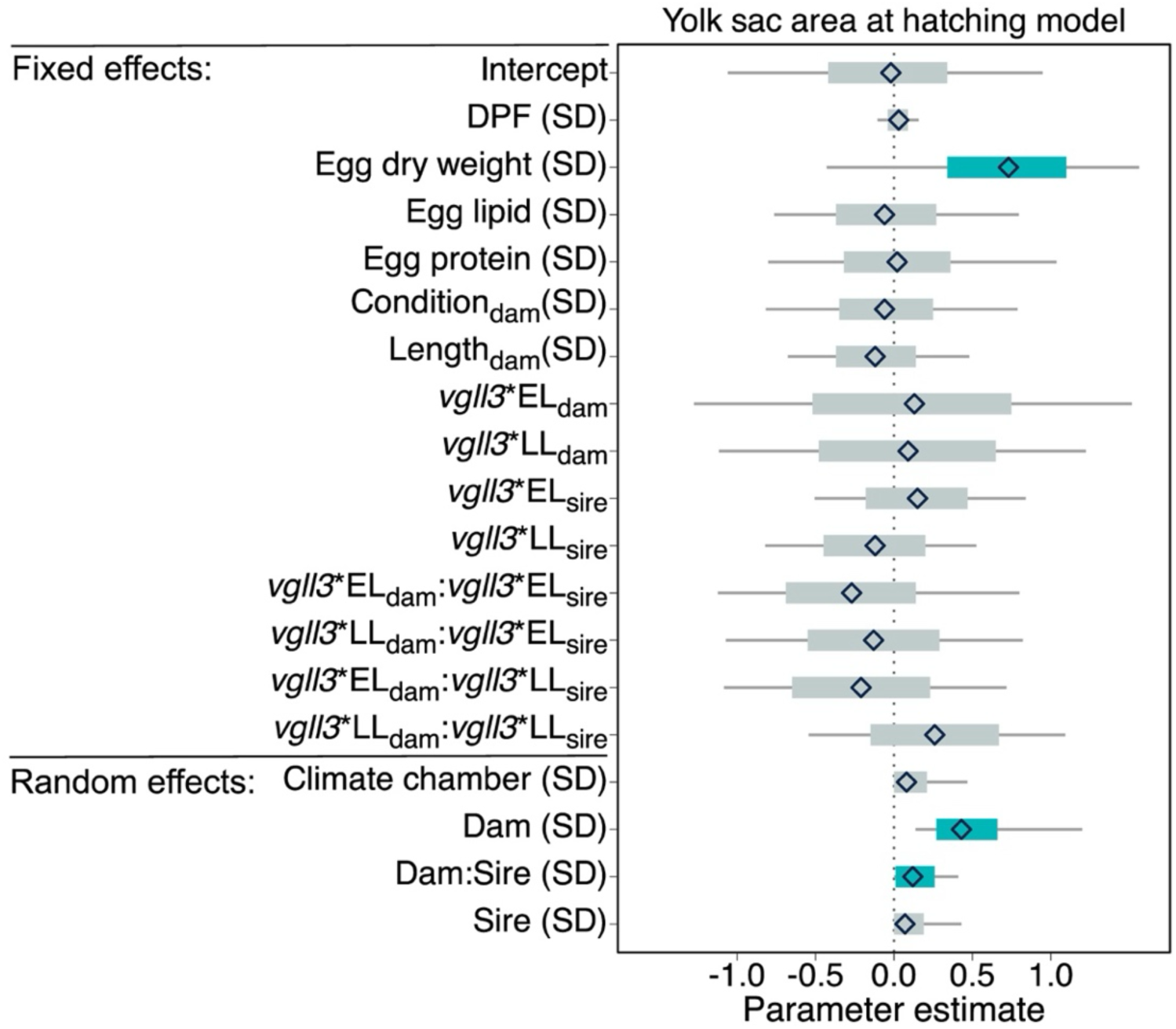
Parameter estimates and 95% credible intervals (CI) for the fixed and random effects from the alevin yolk sac area at hatching model. All continuous variables were mean-centred and standard deviation -scaled prior to fitting the model. Thus, the model estimates indicate the effect of increasing or decreasing the variable by one SD. For the parental *vgll3* genotypes (*vgll3*_dam_, *vgll3*_sire_), the contrast level is the EE genotype. Random effect standard deviations indicate the magnitude of variation between climate chambers and between parents (Dam and Sire) and families (Dam:Sire). The diamonds indicate the mean parameter estimate calculated from 18,000 posterior samples, the thick bars are the 95% CIs, and the thin bars are the 100% CIs. The 95% CI bars are coloured blue if the interval does not contain zero.

**Figure S9.**
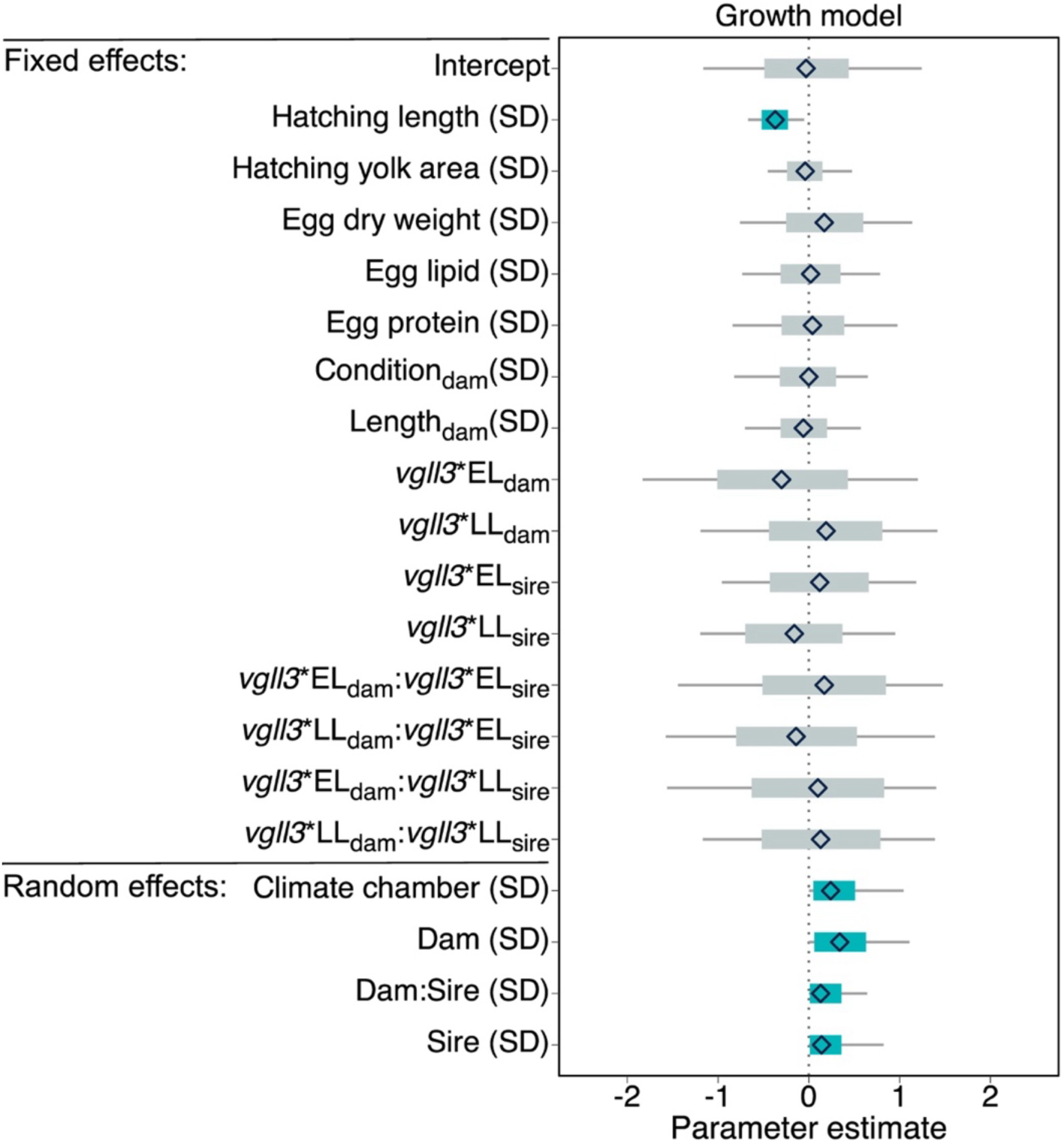
Parameter estimates and 95% credible intervals (CI) for the fixed and random effects from the alevin growth model. All continuous variables were mean-centred and standard deviation -scaled prior to fitting the model. Thus, the model estimates indicate the effect of increasing or decreasing the variable by one SD. For the parental *vgll3* genotypes (*vgll3*_dam_, *vgll3*_sire_), the contrast level is the EE genotype. Random effect standard deviations indicate the magnitude of variation between climate chambers and between parents (Dam and Sire) and families (Dam:Sire). The diamonds indicate the mean parameter estimate calculated from 18,000 posterior samples, the thick bars are the 95% CIs, and the thin bars are the 100% CIs. The 95% CI bars are coloured blue if the interval does not contain zero.

**Figure S10.**
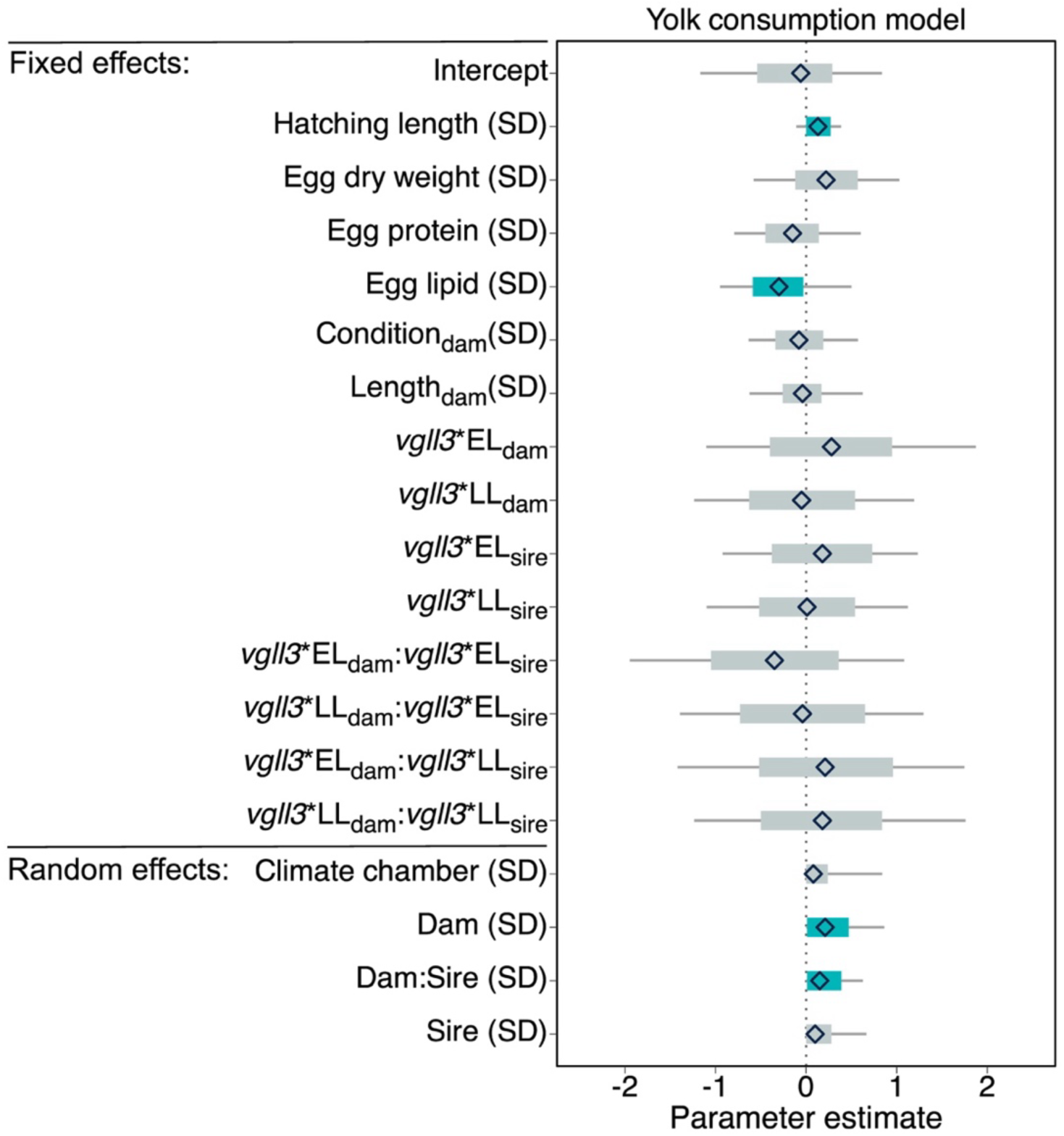
Parameter estimates and 95% credible intervals (CI) for the fixed and random effects from the alevin yolk consumption model. All continuous variables were mean-centred and standard deviation - scaled prior to fitting the model. Thus, the model estimates indicate the effect of increasing or decreasing the variable by one SD. For the parental *vgll3* genotypes (*vgll3*_dam_, *vgll3*_sire_), the contrast level is the EE genotype. Random effect standard deviations indicate the magnitude of variation between climate chambers and between parents (Dam and Sire) and families (Dam:Sire). The diamonds indicate the mean parameter estimate calculated from 18,000 posterior samples, the thick bars are the 95% CIs, and the thin bars are the 100% CIs. The 95% CI bars are coloured blue if the interval does not contain zero.

**Figure S11.**
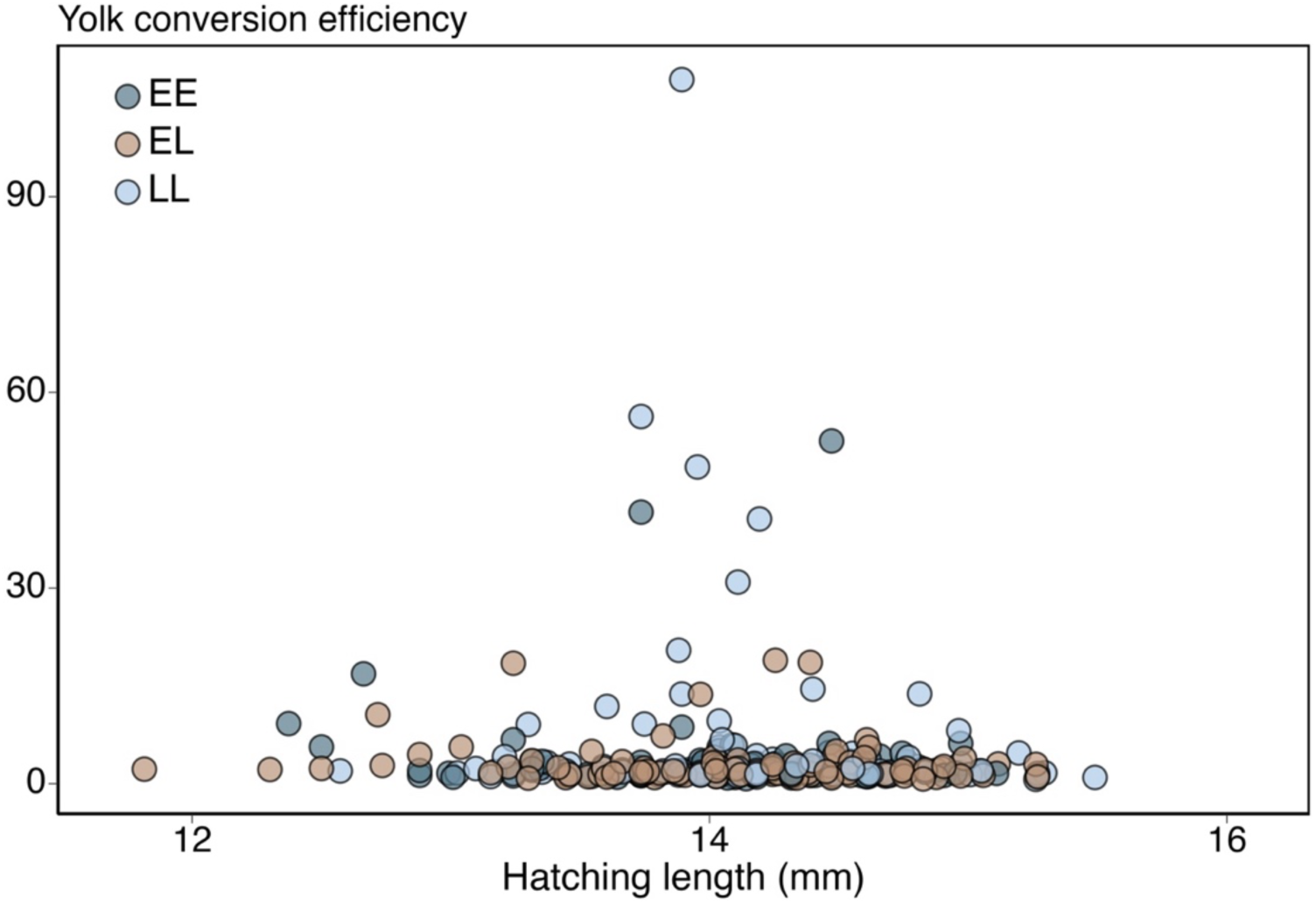
Alevin yolk conversion efficiency between hatching and 81 dpf plotted against hatching length (in mm). The points are coloured according to the maternal *vgll3* genotype (dark blue = EE, light brown = EL, light blue = LL).

**Table S1.** Models used to analyse maternal, egg and alevin traits. Fixed predictors are indicated by their full variable name and random model terms are indicated with a V and subscript. e is the model error term.

| Response variable | Model formulation |
| --- | --- |
| *Maternal phenotype* |  |
| Body length | Body length = $vgll3_{dam} + e$ |
| Body condition | Body condition = $vgll3_{dam} + e$ |
| Fecundity | Fecundity = $vgll3_{dam} + condition_{dam} + length_{dam} + e$ |
| *Egg traits* |  |
| Egg dry weight | Egg dry weight = $vgll3_{dam} + condition_{dam} + length_{dam} + e$ |
| Egg lipid content | Egg lipid = egg dry weight + $vgll3_{dam} + condition_{dam} + length_{dam} + e$ |
| Egg protein content | Egg protein = $vgll3_{dam} + condition_{dam} + length_{dam} + e$ |
| *Alevin traits* |  |
| Family-level alevin survival | Survival = $vgll3_{dam} + vgll3_{sire} + vgll3_{dam} \times vgll3_{sire} + V_{dam} + V_{sire} + V_{dam:sire} + e$ |
| Hatching length | Length<sub>hatching</sub> = DPF + egg dry weight + egg lipid + egg protein + condition<sub>dam</sub> + length<sub>dam</sub> + $vgll3_{dam} + vgll3_{sire} + vgll3_{dam} \times vgll3_{sire} + V_{chamber} + V_{dam} + V_{sire} + V_{dam:sire} + e$ |
| Yolk sac size at hatching | Yolk<sub>hatching</sub> = DPF + egg dry weight + egg lipid + egg protein + condition<sub>dam</sub> + length<sub>dam</sub> + $vgll3_{dam} + vgll3_{sire} + vgll3_{dam} \times vgll3_{sire} + V_{chamber} + V_{dam} + V_{sire} + V_{dam:sire} + e$ |
| Growth | Growth = yolk<sub>hatching</sub> + length<sub>hatching</sub> + egg dry weight + egg lipid + egg protein + condition<sub>dam</sub> + length<sub>dam</sub> + $vgll3_{dam} + vgll3_{sire} + vgll3_{dam} \times vgll3_{sire} + V_{chamber} + V_{dam} + V_{sire} + V_{dam:sire} + e$ |
| Yolk consumption rate (YCR) | YCR = length<sub>hatching</sub> + egg dry weight + egg lipid + egg protein + condition<sub>dam</sub> + length<sub>dam</sub> + $vgll3_{dam} + vgll3_{sire} + vgll3_{dam} \times vgll3_{sire} + V_{chamber} + V_{dam} + V_{sire} + V_{dam:sire} + e$ |
| Yolk conversion efficiency (YCE) | YCE = length<sub>hatching</sub> + egg dry weight + egg lipid + egg protein + condition<sub>dam</sub> + length<sub>dam</sub> + $vgll3_{dam} + vgll3_{sire} + vgll3_{dam} \times vgll3_{sire} + V_{chamber} + V_{dam} + V_{sire} + V_{dam:sire} + e$ |

**Table S2.** Results from the female body length model. Estimate and Est.Error represent the parameter estimate and its error, respectively. 95% CI (lower) and (upper) are the limits of the 95% credible interval and R-hat, Bulk_ESS, and Tail_ESS (ESS = Estimated sample size) describe model convergence and efficiency.

| Fixed effects | Estimate | Est.Error | 95% CI (lower) | 95% CI (upper) | R-hat | Bulk_ESS | Tail_ESS |
| --- | --- | --- | --- | --- | --- | --- | --- |
| Intercept | -0.19 | 0.27 | -0.70 | 0.33 | 1 | 13924 | 12712 |
| $vgll3_{dam} * EL$ | 0.20 | 0.38 | -0.56 | 0.95 | 1 | 14925 | 12183 |
| $vgll3_{dam} * LL$ | 0.36 | 0.36 | -0.37 | 1.07 | 1 | 15751 | 13888 |

**Table S3.** Results from the female body condition model. Estimate and Est.Error represent the parameter estimate and its error, respectively. 95% CI (lower) and (upper) are the limits of the 95% credible interval and R-hat, Bulk_ESS, and Tail_ESS (ESS = Estimated sample size) describe model convergence and efficiency.

| Fixed effects | Estimate | Est.Error | 95% CI<br>(lower) | 95% CI<br>(upper) | R-hat | Bulk_ESS | Tail_ESS |
| --- | --- | --- | --- | --- | --- | --- | --- |
| Intercept | -0.14 | 0.27 | -0.67 | 0.4 | 1 | 16573 | 11404 |
| *vgll3<sub>dam</sub>* *EL | 0.29 | 0.39 | -0.48 | 1.05 | 1 | 16528 | 13235 |
| *vgll3<sub>dam</sub>* *LL | 0.13 | 0.37 | -0.6 | 0.86 | 1 | 16754 | 12987 |

**Table S4.** Results of the fecundity model. Estimate and Est.Error represent the parameter estimate and its error, respectively. 95% CI (lower) and (upper) are the limits of the 95% credible interval and R-hat, Bulk_ESS, and Tail_ESS (ESS = Estimated sample size) describe model convergence and efficiency. Bolded rows indicate model parameters with 95% CI that does not overlap with zero.

| Fixed effects | Estimate | Est.Error | 95% CI<br>(lower) | 95% CI<br>(upper) | R-hat | Bulk_ESS | Tail_ESS |
| --- | --- | --- | --- | --- | --- | --- | --- |
| Intercept | 0.01 | 0.13 | -0.25 | 0.28 | 1 | 18025 | 13252 |
| *vgll3<sub>dam</sub>* *EL | -0.03 | 0.2 | -0.42 | 0.36 | 1 | 19202 | 13801 |
| *vgll3<sub>dam</sub>* *LL | -0.01 | 0.19 | -0.37 | 0.36 | 1 | 18360 | 14429 |
| **Condition (SD)** | **0.3** | **0.08** | **0.13** | **0.46** | **1** | **20866** | **12606** |
| **Length (SD)** | **0.89** | **0.08** | **0.73** | **1.05** | **1** | **20916** | **13829** |

**Table S5.**
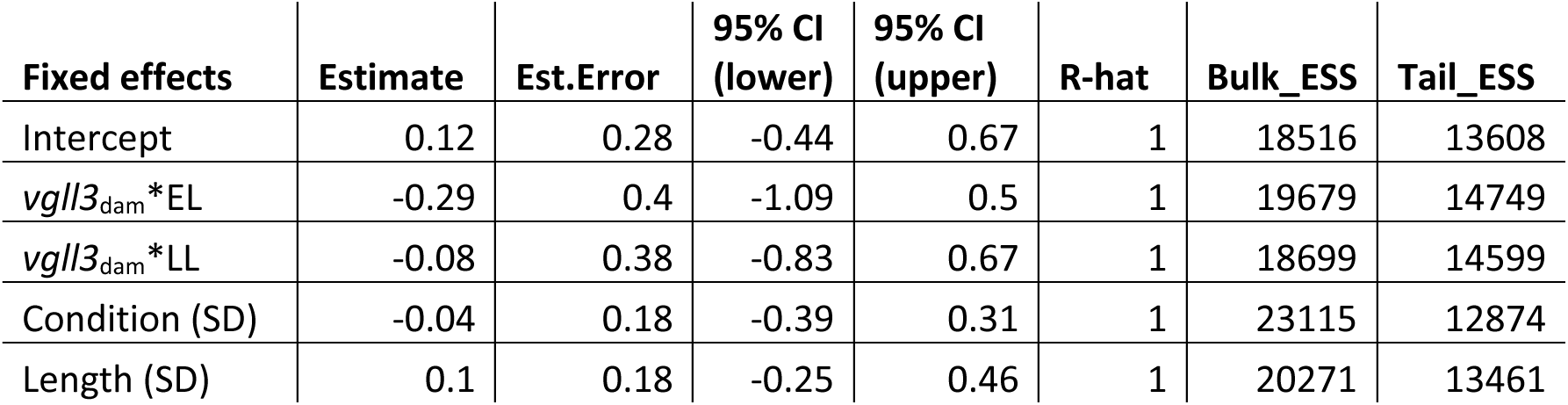
Results of the egg dry weight model. Estimate and Est.Error represent the parameter estimate and its error, respectively. 95% CI (lower) and (upper) are the limits of the 95% credible interval and R-hat, Bulk_ESS, and Tail_ESS (ESS = Estimated sample size) describe model convergence and efficiency.

**Table S6.**
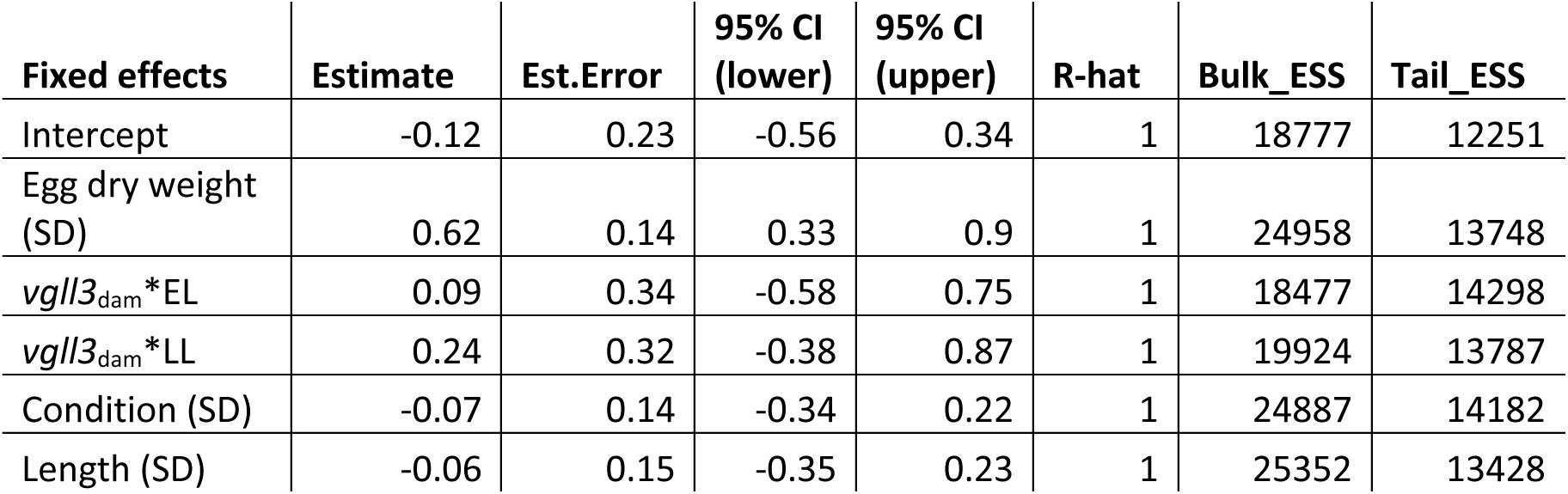
Results of the egg lipid model. Estimate and Est.Error represent the parameter estimate and its error, respectively. 95% CI (lower) and (upper) are the limits of the 95% credible interval and R-hat, Bulk_ESS, and Tail_ESS (ESS = Estimated sample size) describe model convergence and efficiency.

**Table S7.** Results of the egg protein model. Estimate and Est.Error represent the parameter estimate and its error, respectively. 95% CI (lower) and (upper) are the limits of the 95% credible interval and R-hat, Bulk_ESS, and Tail_ESS (ESS = Estimated sample size) describe model convergence and efficiency.

| Fixed effects | Estimate | Est.Error | 95% CI (lower) | 95% CI (upper) | R-hat | Bulk_ESS | Tail_ESS |
| --- | --- | --- | --- | --- | --- | --- | --- |
| Intercept | -0.14 | 0.18 | -0.5 | 0.22 | 1 | 19090 | 13286 |
| *vgll3<sub>dam</sub>* *EL | -0.12 | 0.27 | -0.65 | 0.41 | 1 | 19616 | 13323 |
| *vgll3<sub>dam</sub>* *LL | 0.08 | 0.25 | -0.41 | 0.57 | 1 | 19651 | 14343 |
| Condition (SD) | -0.31 | 0.12 | -0.55 | -0.08 | 1 | 19908 | 12994 |
| Length (SD) | -0.41 | 0.11 | -0.64 | -0.19 | 1 | 20713 | 13112 |

**Table S8.** Results of the alevin survival model. Estimate and Est.Error represent the parameter estimate and its error, respectively. 95% CI (lower) and (upper) are the limits of the 95% credible interval and R-hat, Bulk_ESS, and Tail_ESS (ESS = Estimated sample size) describe model convergence and efficiency.

| **Fixed effects:** | **Estimate** | **Est.Error** | **95% CI**<br>**(lower)** | **95% CI**<br>**(upper)** | **Rhat** | **Bulk_ESS** | **Tail_ESS** |
| --- | --- | --- | --- | --- | --- | --- | --- |
| Intercept | -0.25 | 0.29 | -0.82 | 0.33 | 1 | 5675 | 8109 |
| *vgll3*<sub>dam</sub> * EL | -0.04 | 0.39 | -0.8 | 0.73 | 1 | 6252 | 8541 |
| *vgll3*<sub>dam</sub> * LL | 0.34 | 0.42 | -0.49 | 1.16 | 1 | 6307 | 8655 |
| ***vgll3*<sub>sire</sub> \* EL** | **0.51** | **0.15** | **0.21** | **0.81** | **1** | **12913** | **13103** |
| ***vgll3*<sub>sire</sub> \* LL** | **0.35** | **0.15** | **0.06** | **0.66** | **1** | **12458** | **13082** |
| *vgll3*<sub>dam</sub> * EL:*vgll3*<sub>sire</sub> * EL | -0.29 | 0.19 | -0.67 | 0.09 | 1 | 14853 | 13995 |
| *vgll3*<sub>dam</sub> * LL:*vgll3*<sub>sire</sub> * EL | -0.35 | 0.21 | -0.75 | 0.06 | 1 | 14687 | 13732 |
| *vgll3*<sub>dam</sub> * EL:*vgll3*<sub>sire</sub> * LL | -0.19 | 0.19 | -0.57 | 0.19 | 1 | 14274 | 13177 |
| *vgll3*<sub>dam</sub> * LL:*vgll3*<sub>sire</sub> * LL | -0.28 | 0.2 | -0.68 | 0.12 | 1 | 14746 | 13783 |
| **Random effects:** |  |  |  |  |  |  |  |
| **Dam (SD)** | **0.98** | **0.14** | **0.75** | **1.28** | **1** | **4970** | **7900** |
| **Sire (SD)** | **0.14** | **0.07** | **0.01** | **0.28** | **1** | **3957** | **4740** |

**Table S9.**
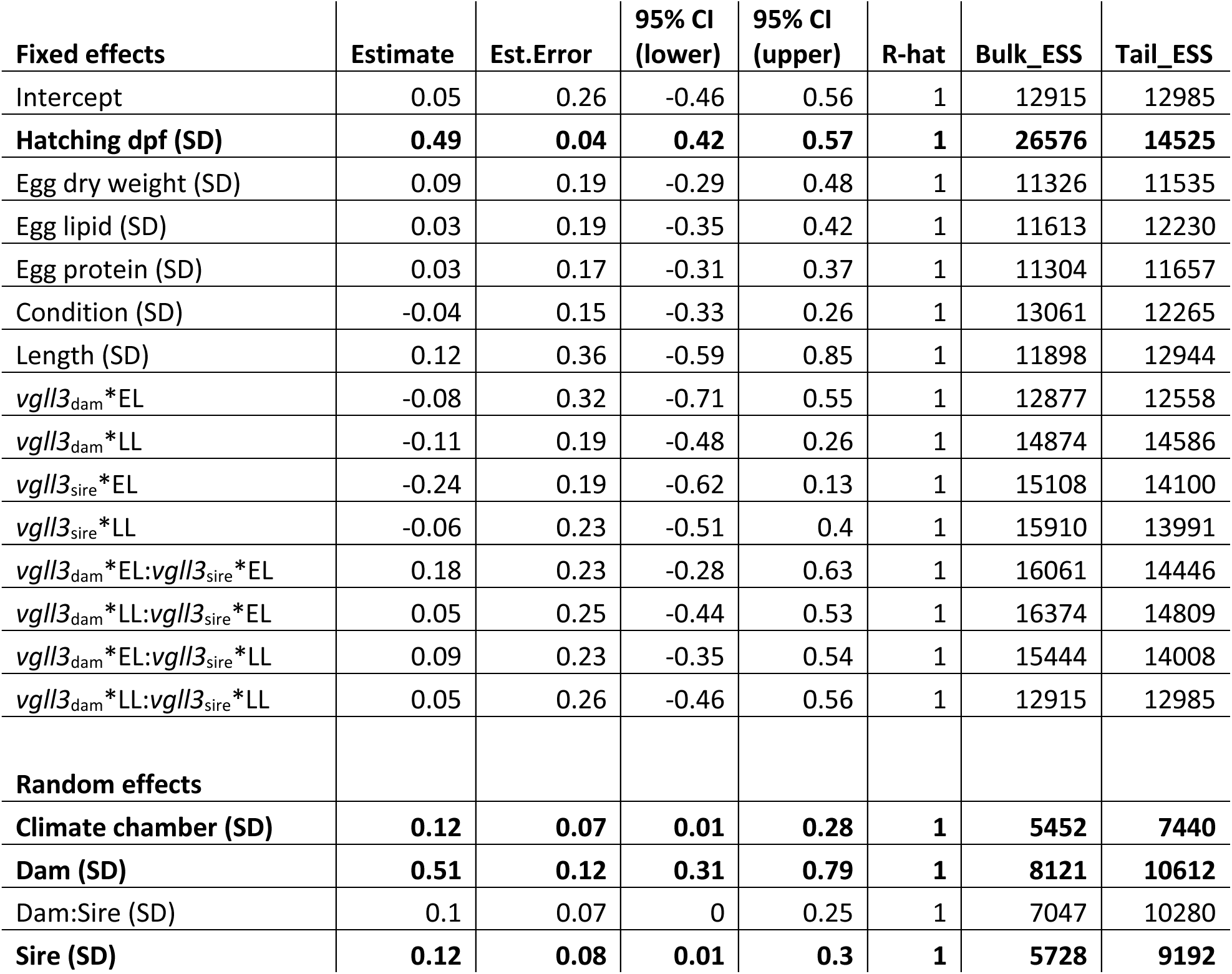
Results of the hatching body length model. Estimate and Est.Error represent the parameter estimate and its error, respectively. 95% CI (lower) and (upper) are the limits of the 95% credible interval and R-hat, Bulk_ESS, and Tail_ESS (ESS = Estimated sample size) describe model convergence and efficiency. Bolded rows indicate model parameters with 95% CI that does not overlap with zero.

| Fixed effects | Estimate | Est.Error | 95% CI (lower) | 95% CI (upper) | R-hat | Bulk_ESS | Tail_ESS |
| --- | --- | --- | --- | --- | --- | --- | --- |
| Intercept | 0.05 | 0.26 | -0.46 | 0.56 | 1 | 12915 | 12985 |
| **Hatching dpf (SD)** | **0.49** | **0.04** | **0.42** | **0.57** | **1** | **26576** | **14525** |
| Egg dry weight (SD) | 0.09 | 0.19 | -0.29 | 0.48 | 1 | 11326 | 11535 |
| Egg lipid (SD) | 0.03 | 0.19 | -0.35 | 0.42 | 1 | 11613 | 12230 |
| Egg protein (SD) | 0.03 | 0.17 | -0.31 | 0.37 | 1 | 11304 | 11657 |
| Condition (SD) | -0.04 | 0.15 | -0.33 | 0.26 | 1 | 13061 | 12265 |
| Length (SD) | 0.12 | 0.36 | -0.59 | 0.85 | 1 | 11898 | 12944 |
| *vgll3*<sub>dam</sub>*EL | -0.08 | 0.32 | -0.71 | 0.55 | 1 | 12877 | 12558 |
| *vgll3*<sub>dam</sub>*LL | -0.11 | 0.19 | -0.48 | 0.26 | 1 | 14874 | 14586 |
| *vgll3*<sub>sire</sub>*EL | -0.24 | 0.19 | -0.62 | 0.13 | 1 | 15108 | 14100 |
| *vgll3*<sub>sire</sub>*LL | -0.06 | 0.23 | -0.51 | 0.4 | 1 | 15910 | 13991 |
| *vgll3*<sub>dam</sub>*EL:*vgll3*<sub>sire</sub>*EL | 0.18 | 0.23 | -0.28 | 0.63 | 1 | 16061 | 14446 |
| *vgll3*<sub>dam</sub>*LL:*vgll3*<sub>sire</sub>*EL | 0.05 | 0.25 | -0.44 | 0.53 | 1 | 16374 | 14809 |
| *vgll3*<sub>dam</sub>*EL:*vgll3*<sub>sire</sub>*LL | 0.09 | 0.23 | -0.35 | 0.54 | 1 | 15444 | 14008 |
| *vgll3*<sub>dam</sub>*LL:*vgll3*<sub>sire</sub>*LL | 0.05 | 0.26 | -0.46 | 0.56 | 1 | 12915 | 12985 |
| **Random effects** |  |  |  |  |  |  |  |
| **Climate chamber (SD)** | **0.12** | **0.07** | **0.01** | **0.28** | **1** | **5452** | **7440** |
| **Dam (SD)** | **0.51** | **0.12** | **0.31** | **0.79** | **1** | **8121** | **10612** |
| Dam:Sire (SD) | 0.1 | 0.07 | 0 | 0.25 | 1 | 7047 | 10280 |
| **Sire (SD)** | **0.12** | **0.08** | **0.01** | **0.3** | **1** | **5728** | **9192** |

**Table S10.** Results of yolk area at hatching model. Estimate and Est.Error represent the parameter estimate and its error, respectively. 95% CI (lower) and (upper) are the limits of the 95% credible interval and R-hat, Bulk_ESS, and Tail_ESS (ESS = Estimated sample size) describe model convergence and efficiency. Bolded rows indicate model parameters with 95% CI that does not overlap with zero.

| **Fixed effects:** | **Estimate** | **Est.Error** | **95% CI (lower)** | **95% CI (upper)** | **R-hat** | **Bulk_ESS** | **Tail_ESS** |
| --- | --- | --- | --- | --- | --- | --- | --- |
| Intercept | -0.04 | 0.23 | -0.49 | 0.41 | 1 | 10191 | 11671 |
| DPF (SD) | 0.03 | 0.03 | -0.04 | 0.09 | 1 | 28011 | 15016 |
| **Egg dry weight (SD)** | **0.73** | **0.2** | **0.34** | **1.1** | **1** | **7603** | **10238** |
| Egg lipid (SD) | -0.06 | 0.16 | -0.37 | 0.27 | 1 | 8440 | 10068 |
| Egg protein (SD) | 0.02 | 0.17 | -0.32 | 0.36 | 1 | 8598 | 10066 |
| Condition (SD) | -0.06 | 0.15 | -0.35 | 0.25 | 1 | 9077 | 9956 |
| Length (SD) | -0.12 | 0.13 | -0.37 | 0.14 | 1 | 9712 | 11097 |
| *vgll3*<sub>dam</sub>*EL | 0.13 | 0.32 | -0.52 | 0.75 | 1 | 9986 | 11912 |
| *vgll3*<sub>dam</sub>*LL | 0.09 | 0.29 | -0.48 | 0.65 | 1 | 10915 | 12136 |
| *vgll3*<sub>sire</sub>*EL | 0.15 | 0.17 | -0.18 | 0.47 | 1 | 11299 | 12496 |
| *vgll3*<sub>sire</sub>*LL | -0.12 | 0.17 | -0.45 | 0.2 | 1 | 11224 | 11645 |
| *vgll3*<sub>dam</sub>*EL:*vgll3*<sub>sire</sub>*EL | -0.27 | 0.21 | -0.69 | 0.14 | 1 | 12685 | 12771 |
| *vgll3*<sub>dam</sub>*LL:*vgll3*<sub>sire</sub>*EL | -0.13 | 0.21 | -0.55 | 0.29 | 1 | 13099 | 12932 |
| *vgll3*<sub>dam</sub>*EL:*vgll3*<sub>sire</sub>*LL | -0.21 | 0.22 | -0.65 | 0.23 | 1 | 12935 | 12726 |
| *vgll3*<sub>dam</sub>*LL:*vgll3*<sub>sire</sub>*LL | 0.26 | 0.21 | -0.15 | 0.67 | 1 | 12496 | 12691 |
| **Random effects:** |  |  |  |  |  |  |  |
| Climate chamber (SD) | 0.08 | 0.05 | 0 | 0.21 | 1 | 5215 | 6574 |
| **Dam (SD)** | **0.43** | **0.1** | **0.27** | **0.66** | **1** | **8627** | **12627** |
| **Dam:Sire (SD)** | **0.12** | **0.07** | **0.01** | **0.26** | **1** | **4400** | **7303** |
| Sire (SD) | 0.07 | 0.05 | 0 | 0.19 | 1 | 9014 | 10869 |

**Table S11.** Results of the alevin growth model. Estimate and Est.Error represent the parameter estimate and its error, respectively. 95% CI (lower) and (upper) are the limits of the 95% credible interval and R-hat, Bulk_ESS, and Tail_ESS (ESS = Estimated sample size) describe model convergence and efficiency. Bolded rows indicate model parameters with 95% CI that does not overlap with zero.

| **Fixed effects:** | **Estimate** | **Est.Error** | **95% CI (lower)** | **95% CI (upper)** | **R-hat** | **Bulk_ESS** | **Tail_ESS** |
| --- | --- | --- | --- | --- | --- | --- | --- |
| Intercept | -0.03 | 0.28 | -0.58 | 0.52 | 1 | 7870 | 11370 |
| **Hatching length (SD)** | **-0.37** | **0.07** | **-0.52** | **-0.23** | **1** | **15874** | **12567** |
| Hatching yolk area (SD) | -0.04 | 0.1 | -0.24 | 0.15 | 1 | 16367 | 14613 |
| Egg dry weight (SD) | 0.17 | 0.21 | -0.25 | 0.6 | 1 | 6857 | 8989 |
| Egg lipid (SD) | 0.02 | 0.17 | -0.31 | 0.35 | 1 | 7451 | 9968 |
| Egg protein (SD) | 0.04 | 0.17 | -0.3 | 0.39 | 1 | 7193 | 9237 |
| Condition (SD) | 0 | 0.16 | -0.32 | 0.3 | 1 | 7326 | 9952 |
| Length (SD) | -0.06 | 0.13 | -0.31 | 0.2 | 1 | 8285 | 10074 |
| *vgll3*<sub>dam</sub>*EL | -0.3 | 0.36 | -1.01 | 0.43 | 1 | 8103 | 10382 |
| *vgll3*<sub>dam</sub>*LL | 0.19 | 0.32 | -0.44 | 0.81 | 1 | 8583 | 11516 |
| *vgll3*<sub>sire</sub>*EL | 0.12 | 0.28 | -0.43 | 0.66 | 1 | 7538 | 10971 |
| *vgll3*<sub>sire</sub>*LL | -0.16 | 0.27 | -0.7 | 0.37 | 1 | 7868 | 11367 |
| *vgll3*<sub>dam</sub>*EL:*vgll3*<sub>sire</sub>*EL | 0.17 | 0.35 | -0.51 | 0.85 | 1 | 8708 | 11073 |
| *vgll3*<sub>dam</sub>*LL:*vgll3*<sub>sire</sub>*EL | -0.14 | 0.34 | -0.8 | 0.53 | 1 | 8686 | 10924 |
| *vgll3*<sub>dam</sub>*EL:*vgll3*<sub>sire</sub>*LL | 0.1 | 0.37 | -0.63 | 0.83 | 1 | 9531 | 13148 |
| *vgll3*<sub>dam</sub>*LL:*vgll3*<sub>sire</sub>*LL | 0.13 | 0.33 | -0.52 | 0.79 | 1 | 8961 | 11877 |
| **Random effects:** |  |  |  |  |  |  |  |
| **Climate chamber (SD)** | **0.24** | **0.12** | **0.05** | **0.51** | **1** | **4230** | **3212** |
| **Dam (SD)** | **0.34** | **0.14** | **0.06** | **0.63** | **1** | **4086** | **3363** |
| **Dam:Sire (SD)** | **0.13** | **0.1** | **0.01** | **0.36** | **1** | **5262** | **8114** |
| **Sire (SD)** | **0.14** | **0.1** | **0.01** | **0.36** | **1** | **4933** | **6022** |

**Table S12.** Results of yolk consumption rate model. Estimate and Est.Error represent the parameter estimate and its error, respectively. 95% CI (lower) and (upper) are the limits of the 95% credible interval and R-hat, Bulk_ESS, and Tail_ESS (ESS = Estimated sample size) describe model convergence and efficiency. Bolded rows indicate model parameters with 95% CI that does not overlap with zero.

| **Fixed effects:** | **Estimate** | **Est.Error** | **95% CI (lower)** | **95% CI (upper)** | **R-hat** | **Bulk_ESS** | **Tail_ESS** |
| --- | --- | --- | --- | --- | --- | --- | --- |
| Intercept | -0.13 | 0.25 | -0.62 | 0.37 | 1 | 9167 | 12077 |
| Hatching length (SD) | 0.13 | 0.07 | 0 | 0.27 | 1 | 31520 | 13593 |
| Egg dry weight (SD) | 0.22 | 0.17 | -0.12 | 0.57 | 1 | 7485 | 9785 |
| Egg protein (SD) | -0.15 | 0.15 | -0.45 | 0.14 | 1 | 7702 | 10321 |
| **Egg lipid (SD)** | **-0.3** | **0.14** | **-0.59** | **-0.03** | **1** | **8752** | **11009** |
| Condition (SD) | -0.08 | 0.13 | -0.34 | 0.19 | 1 | 8417 | 10893 |
| Length (SD) | -0.04 | 0.11 | -0.26 | 0.17 | 1 | 9119 | 10668 |
| *vgll3*<sub>dam</sub>*EL | 0.28 | 0.34 | -0.4 | 0.95 | 1 | 9515 | 12154 |
| *vgll3*<sub>dam</sub>*LL | -0.05 | 0.3 | -0.63 | 0.54 | 1 | 9891 | 11639 |
| *vgll3*<sub>sire</sub>*EL | 0.18 | 0.28 | -0.38 | 0.73 | 1 | 10185 | 13026 |
| *vgll3*<sub>sire</sub>*LL | 0.01 | 0.27 | -0.52 | 0.54 | 1 | 9710 | 11377 |
| *vgll3*<sub>dam</sub>*EL:*vgll3*<sub>sire</sub>*EL | -0.35 | 0.36 | -1.05 | 0.36 | 1 | 9900 | 12153 |
| *vgll3*<sub>dam</sub>*LL:*vgll3*<sub>sire</sub>*EL | -0.04 | 0.36 | -0.73 | 0.65 | 1 | 11504 | 12733 |
| *vgll3*<sub>dam</sub>*EL:*vgll3*<sub>sire</sub>*LL | 0.21 | 0.38 | -0.52 | 0.96 | 1 | 11092 | 13191 |
| *vgll3*<sub>dam</sub>*LL:*vgll3*<sub>sire</sub>*LL | 0.18 | 0.34 | -0.5 | 0.84 | 1 | 10865 | 12886 |
| **Random effects:** |  |  |  |  |  |  |  |
| Climate chamber (SD) | 0.08 | 0.06 | 0 | 0.24 | 1 | 10609 | 9674 |
| **Dam (SD)** | **0.21** | **0.12** | **0.01** | **0.47** | **1** | **4541** | **6297** |
| **Dam:Sire (SD)** | **0.15** | **0.1** | **0.01** | **0.39** | **1** | **5427** | **7821** |
| Sire (SD) | 0.1 | 0.08 | 0 | 0.28 | 1 | 9300 | 10151 |

**Table S13.** Results of yolk conversion efficiency model. Estimate and Est.Error represent the parameter estimate and its error, respectively. 95% CI (lower) and (upper) are the limits of the 95% credible interval and R-hat, Bulk_ESS, and Tail_ESS (ESS = Estimated sample size) describe model convergence and efficiency.

| **Fixed effects:** | **Estimate** | **Est.Error** | **95% CI (lower)** | **95% CI (upper)** | **R-hat** | **Bulk_ESS** | **Tail_ESS** |
| --- | --- | --- | --- | --- | --- | --- | --- |
| Intercept | -0.23 | 0.04 | -0.3 | -0.15 | 1 | 6559 | 8447 |
| Hatching dpf (SD) | -0.01 | 0.01 | -0.03 | 0 | 1 | 27365 | 15195 |
| Egg dry weight (SD) | 0.01 | 0.02 | -0.04 | 0.05 | 1 | 9252 | 10933 |
| Egg lipid (SD) | 0.02 | 0.02 | -0.01 | 0.06 | 1 | 10977 | 10707 |
| Egg protein (SD) | 0 | 0.02 | -0.03 | 0.03 | 1 | 9274 | 11079 |
| Condition (SD) | -0.01 | 0.02 | -0.04 | 0.02 | 1 | 10471 | 12744 |
| Length (SD) | 0 | 0.01 | -0.03 | 0.02 | 1 | 10952 | 12093 |
| *vgll3*<sub>dam</sub>*EL | -0.02 | 0.05 | -0.12 | 0.07 | 1 | 7067 | 9827 |
| *vgll3*<sub>dam</sub>*LL | -0.01 | 0.04 | -0.09 | 0.08 | 1 | 7667 | 9426 |
| *vgll3*<sub>sire</sub>*EL | -0.04 | 0.04 | -0.13 | 0.04 | 1 | 7469 | 10296 |
| *vgll3*<sub>sire</sub>*LL | -0.01 | 0.04 | -0.1 | 0.07 | 1 | 7077 | 10688 |
| *vgll3*<sub>dam</sub>*EL:*vgll3*<sub>sire</sub>*EL | 0.05 | 0.05 | -0.05 | 0.15 | 1 | 8163 | 10863 |
| *vgll3*<sub>dam</sub>*LL:*vgll3*<sub>sire</sub>*EL | 0.01 | 0.05 | -0.09 | 0.1 | 1 | 8241 | 10352 |
| *vgll3*<sub>dam</sub>*EL:*vgll3*<sub>sire</sub>*LL | -0.01 | 0.05 | -0.12 | 0.1 | 1 | 7894 | 11479 |
| *vgll3*<sub>dam</sub>*LL:*vgll3*<sub>sire</sub>*LL | 0.02 | 0.05 | -0.08 | 0.12 | 1 | 8149 | 11814 |
| **Random effects:** |  |  |  |  |  |  |  |
| Climate chamber (SD) | 0.01 | 0.01 | 0 | 0.03 | 1 | 10241 | 8991 |
| Dam (SD) | 0.02 | 0.01 | 0 | 0.05 | 1 | 4886 | 6439 |
| Dam:Sire (SD) | 0.02 | 0.01 | 0 | 0.04 | 1 | 4787 | 7173 |
| Sire (SD) | 0.02 | 0.01 | 0 | 0.04 | 1 | 5440 | 8363 |

